# Observing concurrent subcellular dynamics in large living tissues

**DOI:** 10.64898/2026.02.25.707857

**Authors:** Charles S Wright, Sanjeev Uthishtran, Laura Z Kreplin, Hetvi R Gandhi, Abhishek Patil, Harrison M York, Samyukta Sita, Samuel A Manning, Elliot Brooks, Guizhi Sun, In-won Lee, Wing Hei Chan, Sara Hlavca, Samuel Crossman, Helen E Abud, Jan Kaslin, Avnika A Ruparelia, Peter D Currie, Kieran F Harvey, Jose M Polo, John Carroll, Senthil Arumugam

**Affiliations:** Department of Anatomy & Developmental Biology, Monash Biomedicine Discovery Institute, Monash University, Melbourne, VIC, Australia; EMBL Australia, Monash University, Melbourne, VIC, Australia; Peter MacCallum Cancer Centre, Melbourne, VIC, Australia; Cabrini Monash University Department of Surgery, Cabrini Hospital, Malvern, Australia; Australian Regenerative Medicine Institute, Monash University, Clayton, VIC, Australia; Centre for Muscle Research, Department of Anatomy and Physiology, University of Melbourne, Melbourne, Victoria, 3010, Australia; Department of Anatomy and Physiology, School of Biomedical Sciences, Faculty of Medicine Dentistry and Health Sciences, University of Melbourne, Melbourne, Victoria, 3010, Australia; Sir Peter MacCallum Department of Oncology, The University of Melbourne, Parkville, VIC, Australia; The Adelaide Centre for Epigenetics and the South Australian Immunogenomics Cancer Institute, Faculty of Health and Medical Sciences, The University of Adelaide, Adelaide, SA, Australia

**Author notes:** Aix Marseille Université, CNRS, INP UMR7051, NeuroCyto, Marseille, France. Equal first authors (Charles S Wright, Sanjeev Uthishtran, Laura Z Kreplin, Hetvi R Gandhi, Abhishek Patil, Harrison M York).

## Abstract

An outstanding question in eukaryotic biology is the mechanistic connection between events occurring at (sub)cellular levels (time scales of milliseconds to minutes) to those at the tissue levels (tens of minutes to months). Deciphering such mechanisms requires imaging approaches capable of simultaneously achieving high spatial and temporal resolutions for large samples over long periods of time. Here, we demonstrate Airy beam-based light sheet microscopy of organelles in tens to hundreds of cells in a few hundred micrometre-wide tissue environments. We achieve a typical resolution of 320 nm over 266 × 266 × 100 μm^3^ volumes at a temporal rate of 0.05 Hz, typically with generally used fluorophores such as Green Fluorescent Protein, over extended periods of time that allow tracking of organelle and protein dynamics. We validated our approach across different length and time scales by imaging mitochondria and endosome dynamics in very large fields of view in zebrafish tissue, molecular assemblies of myosin as gastrulation proceeds in *Drosophila* embryos, 3D mitochondrial streaming in mouse oocytes, pressure-driven motility and protrusions in amoebae, mitochondrial dynamics in cancer spheroids, 5 -colour fast imaging in iBlastoids, and endosomal dynamics in single cells. Through these model systems, we demonstrate the versatility of Airy beam light sheet microscopy to image large tissues at unprecedented high resolution; to capture dynamics in photosensitive, delicate samples; and to screen 3D samples. We anticipate that our Airy beam-based approach will represent a pivotal advance in cellular biology—especially developmental biology—as it provides, for the first time, true subcellular resolution over large imaging volumes with high temporal resolution.

## Introduction

Multiscale measurements are critical to addressing the fundamental question of how molecular and cellular events give rise to emergent tissue-level behaviours. This problem is particularly evident in the context of animal development, which involves processes that span the extremes of biological length and time scales and integrate genetic, biochemical, and mechanical information ( *1*). Gene regulation, intra- and intercellular signalling, organellar dynamics, and cell shape changes and movements (taking place over milliseconds to minutes) drive tissue development and sculpting (occurring over tens of minutes to months), which in turn feeds back to modulate lower-level components. This interplay between processes occurring at multiple, hierarchical levels of organisation—linking transcriptional programs to mechanistic execution through signalling and mechanochemical pathways, resulting in macroscopic morphogenetic processes—is thus crucial for coordinating spatial events and generating the temporal patterns required for robust development.

However, it is also one of the least explored facets of developmental biology, due largely to measurement challenges. Connecting stochastic components at the molecular and organellar levels (small length scales, fast time scales) to emergent behaviours at higher levels of organisation (large length scales, slow time scales) requires an identification of the cross-scale interactions of patterns and processes that is only obtainable by observing the relevant dynamics within living organisms. Thus, a major bottleneck in understanding mechanisms that operate across multiple levels of organisation has been the ability to make simultaneous measurements across the corresponding spatial and temporal scales.

Fluorescence microscopy has yielded tremendous advances in our understanding of biological processes whose details may be captured within the spatial and temporal windows compatible with specific imaging modalities. At the molecular level, specialised approaches, including super-resolution microscopy, have proven highly effective in elucidating the dynamics and organisation of interacting molecules within living cells (*2*). At the subcellular level, various imaging technologies have advanced our understanding of biochemical processes that regulate organelle dynamics and their transport, organisation, and regulation (*3, 4*). At the levels of cells and tissues, *in toto* imaging approaches typically leverage light sheet-based modalities, which illuminate and image thin planes of light scanned rapidly through a 3D sample. These studies have enabled reconstruction of cell lineages, primarily within the model organisms *Drosophila* (*5*), zebrafish (*6*), mouse embryo (*7, 8*), arthropod limbs (*9*) and body-wide circuits of cellular activities and pulsating waves of calcium across tissues (*10*). However, the bridge between molecular interactions, organelle dynamics within single cells, and morphogenesis within tissues, remains poorly explored. For example, a challenging question in the field is the connection between rapid intracellular organelle dynamics, such as endosomes and mitochondria, within the context of developing tissue spanning many cells. Light sheet-based approaches can capture fast organelle dynamics at whole-cell volumes for single cells (*3*), and more recently, adaptive optics-based lattice light-sheet microscopy (LLSM) has been used to observe organelle dynamics in living tissues and organoids (*11*). However, given the very limited field of view (FoV), a large volume can be achieved only by tiling multiple sub volumes, which significantly reduces temporal resolution and requires stitching. Within light sheet-based approaches, the size of the lateral FoV is largely determined by the choice of beam. Motivated by the need to study fast developmental processes, including mapping organelle dynamics and morphological cellular and morphogenetic tissue changes simultaneously, and the lack of a technique capable of imaging at the requisite spatial and temporal scales, we explored beams that could offer the versatility and uniform excitation across the FoV needed to permit tiling-free capture of large volumes.

Light-sheet approaches using Bessel beams, which have cylindrically symmetric profiles, have successfully imaged subcellular dynamics in living cells (*12*), both using single beams as well as multiple interfering beams as in the case of LLSM (*13*). Like Bessel beams, Airy beams are non-diffracting and enable optical sectioning; however, their asymmetric excitation pattern can result in enhanced contrast (*14*). While low numerical aperture (NA) systems using Airy beams have been demonstrated for large tissues at low resolution (*14*), imaging at subcellular resolution that combines Airy beams with high-NA optics in an optimised fashion to span large FOVs with expanded spatiotemporal coverage in large living tissues has yet to be shown. Key characteristics and benchmarks of existing light-sheet approaches are summarised in Supplementary table T1.

Here, we report a light-sheet microscope based on versatile Airy beams that are optimised for the NA of the excitation objective and the FoV relative to the camera chip size, in combination with a high-NA water immersion collection objective. This larger, optimised FoV translates to enhanced temporal resolution (speed) for large tissues while maximising the spatial resolution required to capture organelles. Consequently, our approach achieves 0.05 Hz temporal resolution over 266 × 266 × 100 μm^3^ at 320 nm resolution. This is a significant improvement compared to previous imaging solutions (Supplementary table T1). To demonstrate the versatility of this approach for biomedical imaging, we imaged mitochondria and endosomes in zebrafish embryos and larvae, mid-gut invagination in *Drosophila* embryos, mitochondrial streaming in mouse oocytes, amoeba motility, and mitochondrial and cytoskeletal dynamics in organoids. We also show that rapid imaging allows high-throughput screening in iBlastoids, and that shorter Airy beams can be used to measure organelle dynamics in cells. Through these examples, we demonstrate the capability of Airy beams in enlarging the available spatiotemporal resolution regime, which opens the door to investigate previously inaccessible biological questions requiring ‘across-the-scale’ imaging and measurements, including mapping protein movements, molecular assembly dynamics, organelle dynamics, and cell motilities across large volumes of living tissues.

## Results

### Versatility of Airy beams to balance field of view versus resolution

To realise an Airy beam light-sheet microscope system, we utilised the previously established geometry of high-NA objective pairings used for Bessel and lattice light-sheet microscopy. We utilised a 20× 0.6 NA Thorlabs objective to deliver excitation light and a 25× 1.1 NA Nikon objective or a 20× 1.0 NA Olympus (Evident) objective for collection of emission light, providing a 266 × 266 μm^2^ or a 332 × 332 μm^2^ FoV, respectively, for a single plane. We used a spatial light modulator (SLM) at the Fourier plane to the sample plane to modulate the incoming beam with a cubic phase, resulting in an Airy beam after the excitation objective (Materials and methods, Supplementary fig. 1). By applying different scale factors, the non-diffractive propagation distance can be modulated to fill the FoV of the camera to distinct extents. We scan this beam across the entire FoV using a galvo mirror. There is no scanning along the direction of propagation of light after the detection objective, but modulation of the Airy beam to different lengths using the SLM offers a choice between a smaller FoV with a higher resolution, or a larger FoV with a lower resolution. Scanning across multiple planes can be performed in three ways (Fig. 1a). First, an ‘XZ - scan’ can be performed where the sample is moved along the principal axis of the collection objective, which is achieved by mounting the stage motor at the same angle as the collection objective with respect to the base of the optical table. This approach abrogates any need for post-acquisition deskew processes. Second, the stage can be moved parallel to the optical table in an ‘X-scan’, permitting imaging of larger areas of the sample by moving them laterally, but requiring deskew. Third, a galvo-electronic tuneable lens (ETL) is used, where the sample is not moved at all, but the beam is scanned across using a galvo mirror, and the focus of the collection objective is shifted in synchrony using the ETL. This approach does not require deskew, but the range of motion is limited compared to the XZ-scan.

**Figure 1.**
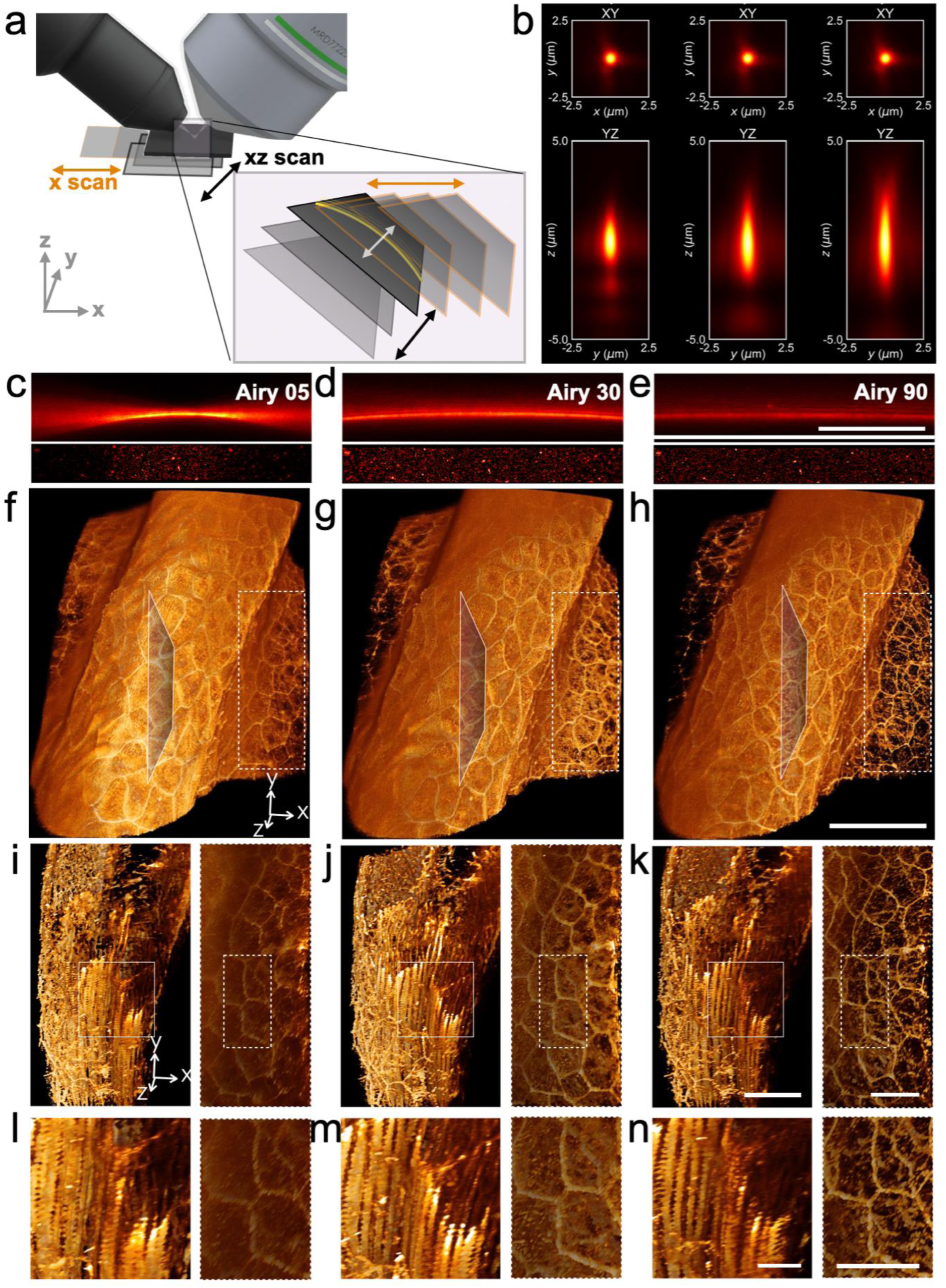
Airy beam light-sheet microscope acquisition modes and beam types. **(a)** Schematic illustrating main acquisition types. The XZ-scan moves along the same axis as the detection objective and is suitable for most samples. The X-scan moves along the same axis as the optical table and is suitable for scans through wide but relatively thinner samples. **(b)** Experimentally measured PSFs of Airy beams corresponding to phase masks of increasing strength (Airy 5, 30, and 90, respectively). *Top*: XY projections. *Bottom*: YZ projections. Images were obtained with 561 nm excitation wavelength and 100 nm diameter TetraSpeck beads. **(c–e)** *Top:* Experimentally measured beam profiles corresponding to the same phase masks as in **b**. *Bottom:* Section through a volume of 100 nm diameter TetraSpeck beads illustrating horizontal variation in intensity and resolution accompanying different Airy beams as in **b**. **(f–n)** Alexa Fluor 647-labelled phalloidin in zebrafish tissue measured with the same Airy beams as in **b**. **(f–h)**The sample spans 266 × 266 × 320 µm^3^. **(i–k)** Each block of insets shows medium zoom cross-section views taken from regions near the centre (*left*), representing the dorsal tail tissue, and the edge (*right*), representing the yolk tissue, of the corresponding images in **f–h**. A further zoom of the marked regions in **i–k** is displayed in **l–n**. Scale bars: **(c–h):** 100 μm, **(i–k):** 25 μm, **(l–n):** 5 μm.

To characterise the resolution and FoV capabilities of our system, we used 100 nm TetraSpeck beads that produce diffraction-limited images. Different Airy beams cover the FoV to different extents (Fig. 1c–e, Supplementary fig. 2a) and produce distinct PSFs (Fig. 1b, Supplementary fig. 2b). As is evident from the PSFs, the compromise for the FoV is the axial resolution: raw images acquired using Airy beams have a characteristic blur in the axial direction, necessitating deconvolution to achieve maximal axial resolution. Here, all images displayed have been deconvolved using PSFs calculated from diffraction-limited images of 100 nm TetraSpeck beads for their respective wavelengths and Airy beams. We then imaged phalloidin-stained zebrafish tissue spanning 266 × 266 × 320 μm^3^ (Fig. 1f–n). As expected, with longer Airy beams, resolution improved at full volume-in-view (ViV). The XZ-scan, in combination with an appropriate Airy beam length, was also able to capture organelle dynamics in single cells. We next captured endosomal motility in single cells; dual-channel images were simultaneously acquired from HeLa cells expressing Rab-GFP and Rab7-mCherry, spanning 103 × 302 × 25 μm^3^ and imaged over 17 min at 5.7 s per volume (Supplementary fig. 3, Supplementary movie 1), demonstrating the versatility of our approach from large tissues to single cells.

We estimated resolution using Fourier ring correlation (*15*) in both single cells with JFX650-labelled mitochondria (Fig. 2a), as well as phalloidin-stained zebrafish embryos (Fig. 2 b,c). While the effective resolution was dependent on the local structure and signal-to-noise, it was on average ∼320 nm and did not degrade with depth for zebrafish, a transparent sample. In both cases, for single cells and tissues, we demonstrate that increasing the beam length resulted in improvement of the ViV, albeit with compromised highest achievable spatial resolution. More importantly, while comparisons on resolutions can be made on many static parameters calculated from inanimate samples such as fluorescent beads, we show in the next sections that live imaging of biological processes could be performed, the metric most pertinent to the biological community ( *16*). See Supplementary table T2 for a summary of all biological samples, imaging parameters, and benchmarking metrics.

**Figure 2.**
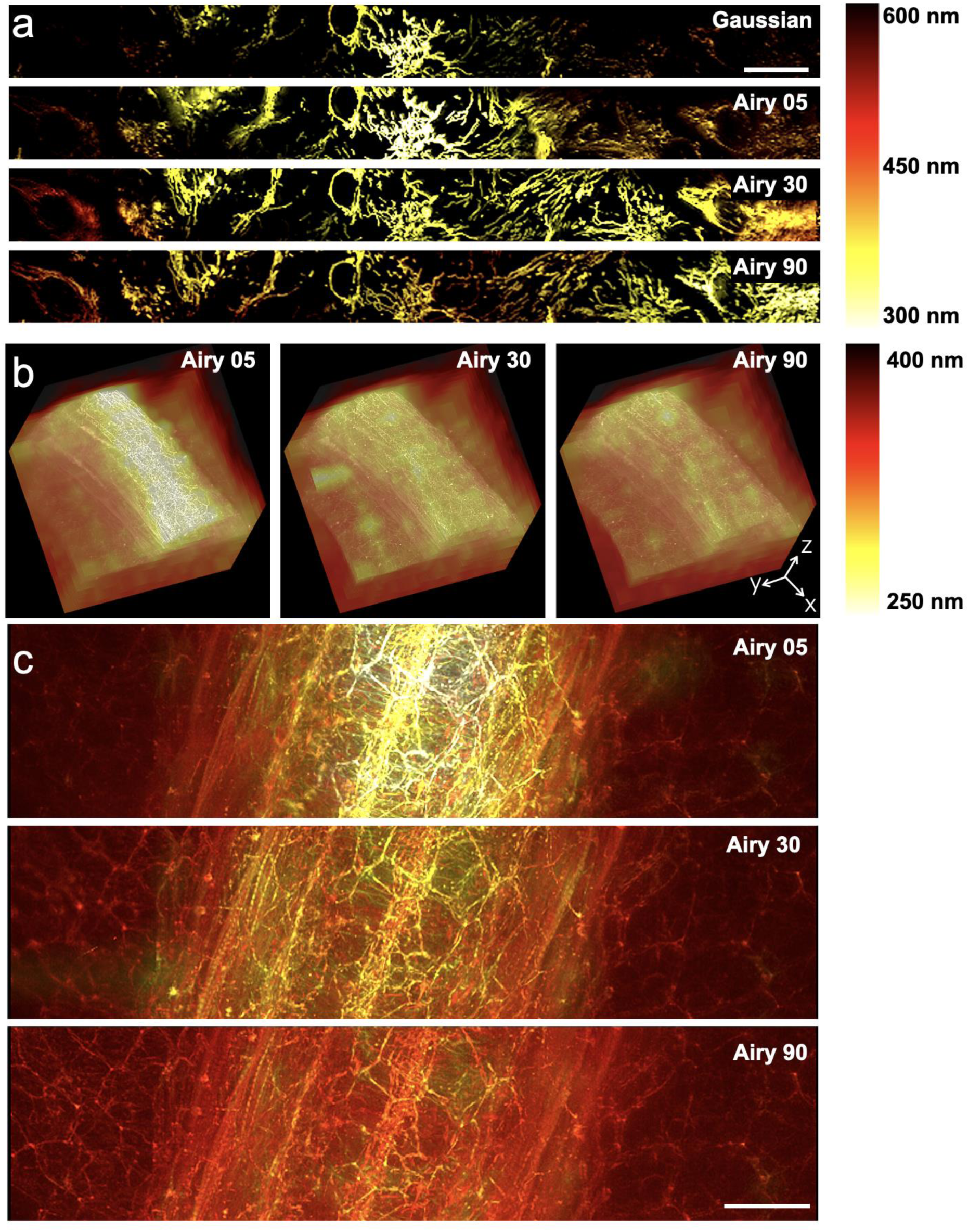
Resolution maps calculated from live samples. **(a)** Scan through a thin sample of confluent HeLa cells with JFX650-labelled mitochondria illustrating the variation in lateral resolution as a function of Airy beam strength across a wide (332 μm) region. The image has been colour-coded according to the calculated resolution. Rows are arranged from top to bottom in order of increasing Airy beam strength. Images were acquired with a 20× 1.0 NA collection objective. **(b–c)** Section of a thick (266 × 266 × 320 µm^3^) volume of fixed zebrafish tissue labelled with phalloidin-Alexa Fluor 647 illustrating the trade-offs between maximum achievable resolution in the centre of the FoV (lower beam strength) and maximal effective resolution across the full FoV (higher beam strength). Images were acquired with a 25× 1.1 NA collection objective. **(b)** Resolution maps, with columns arranged from left to right in order of increasing Airy beam strength. **(c)** Section along the length of the Airy beam, where images have been colour-coded according to the calculated resolution. Note that the difference in the observed limits of the resolution between **a** and **b–c** is largely due to the collection objective. Scale bars: **(a,c):** 50 μm.

### High spatiotemporal, large-volume, dynamic measurements in zebrafish

To demonstrate our approach in measuring dynamic events across the entire volume of imaging (VoI) in living tissues, we imaged GFP-labelled mitochondria and mCherry-labelled endoplasmic reticulum (ER) in the skeletal muscle of transgenic *Tg(actc1b:mito-GFP)^uom407Tg^* and *Tg(actc1b:ER-mCherry)^uom408Tg^* double-positive zebrafish larvae at 3 days post fertilisation (dpf). Maximising the illumination across the entire FoV, and using the XZ-scan, we could image a volume of 266 × 266 × 40 μm^3^ with a time resolution of 19 s per volume or at a rate of 0.05 Hz for two colours (Fig. 3a, Supplementary movie 2). The uniform resolution across the entire volume allows ensemble quantification by precise segmentation (Fig. 3b, Supplementary movie 3) and analysis of the orientation of mitochondria and ER in individual myotomes (Fig. 3c). Furthermore, the high time resolution enabled visualisation of mitochondrial dynamics at multiple locations at the same time (green, magenta, and cyan highlighted regions, Fig. 3d; Supplementary movie 4) revealing fusion and fission events (Fig. 3e; colours correspond to regions highlighted in Fig. 3d). The consistent resolution across the volume also enables morphometric analysis of nuclei or plasma membranes with high accuracy (Supplementary fig. 4,5).

**Figure 3.**
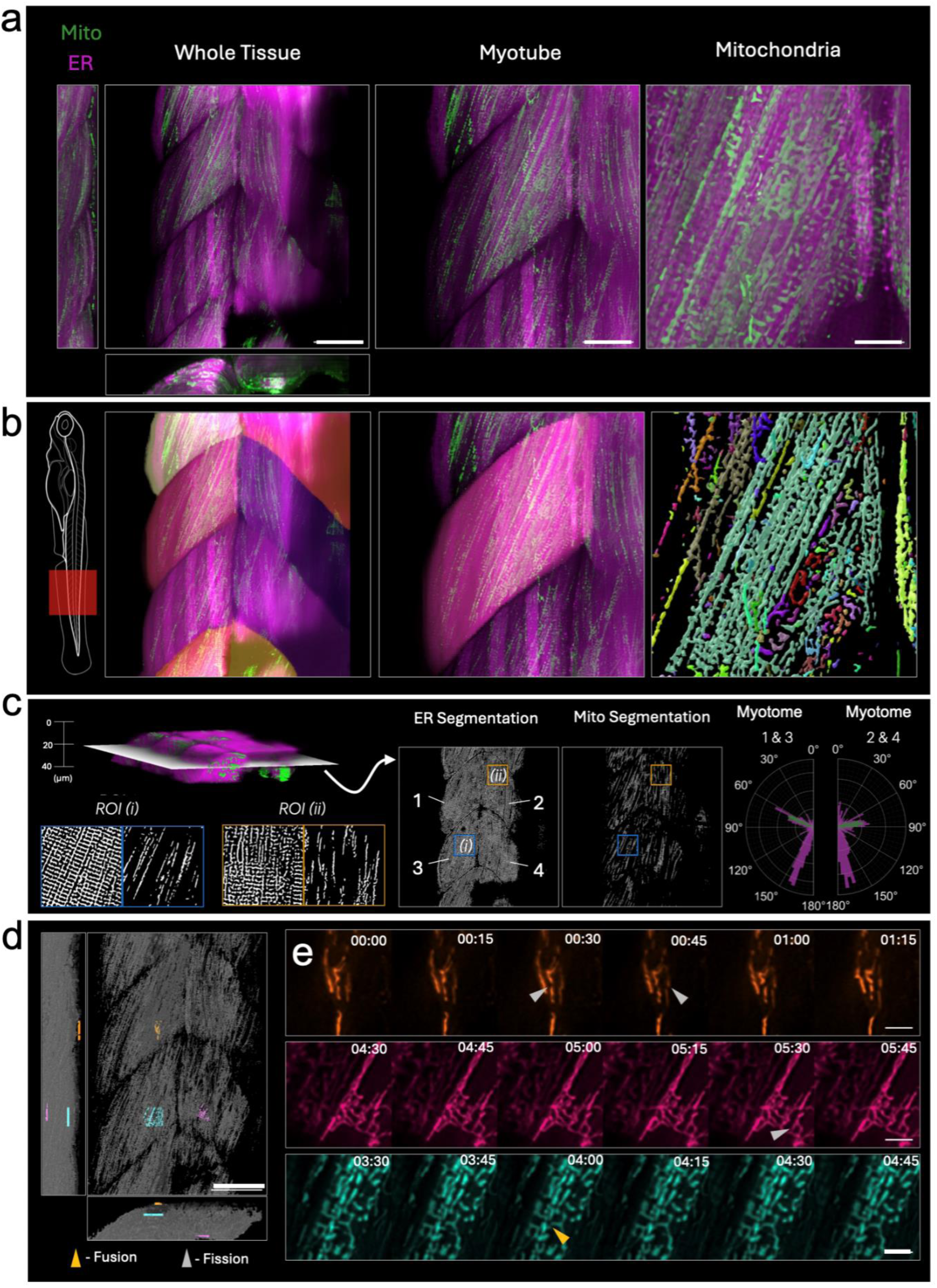
Organelle dynamics in large volumes of zebrafish tissue. **(a)** Raw images of skeletal muscle tissue of a 3 dpf transgenic *Tg(actc1b:mito-GFP)^uom407Tg^* and *Tg(actc1b:ER-mCherry)^uom408T^*^g^ double positive larva at progressively higher zooms, showcasing different scales of organisation from the whole tissue to individual myotomes and mitochondria. **(b)** Schematic of imaged region of zebrafish larva (*left*). Corresponding 3D segmentations of the regions displayed in **a**, depicting the whole tissue, individual myotomes, and mitochondria. **(c)** Cross-sectional slice extracted from the middle of the volume of a representative tissue section, displaying both the endoplasmic reticulum (ER) and mitochondrial channels, alongside their respective segmentations. Two 36 μm × 36 μm zoomed-in regions of interest (ROIs), indicated by blue and orange boxes, highlight distinct sub-regions within the segmented volume. Semi-polar plots illustrating the orientation of ER banding (magenta) and mitochondrial networks (green). The left plot presents data from myotomes 1 and 3. The right plot displays data from myotomes 2 and 4. **(d)** Segmentation of an entire muscle tissue volume with three independently selected ROIs, colour-coded orange, magenta, and cyan. **(e)** A montage of deconvolved raw time-series data from the corresponding colour-coded ROIs in **d** displaying mitochondrial dynamics. Mitochondrial fission (grey arrows) and fusion (yellow arrows) events are highlighted. Scale bars: **(a,d):** 50 μm (whole tissue), 30 μm (myotomes), and 10 μm (mitochondria); **(d):** 50 μm; **(e)**: 5 μm.

To demonstrate concurrent, continuous tracking across a large volume at high temporal resolution, we imaged mKate2-rab5ab-positive endosomes across 266 × 266 × 60 μm^3^ in the tailbud tissue of a 14-somite stage zebrafish embryo expressing mKate2-rab5ab with a time resolution of 5.9 s per volume (Fig. 4a–d; Supplementary movies 5,6). Using a custom code for detecting and volumetric tracking of endosomes (*17*), we were able to capture endosomal trajectories within individual cells across the VoI and characterise the motility of endosomes both within and between cells of a large cross-section of developing tissue (Fig. 4e). We also imaged a zebrafish embryo expressing mKate2-rab5ab and membrane-mNeonGreen to visualise both endosomal motility as well as cell boundaries (Fig. 4f–h).

**Figure 4.**
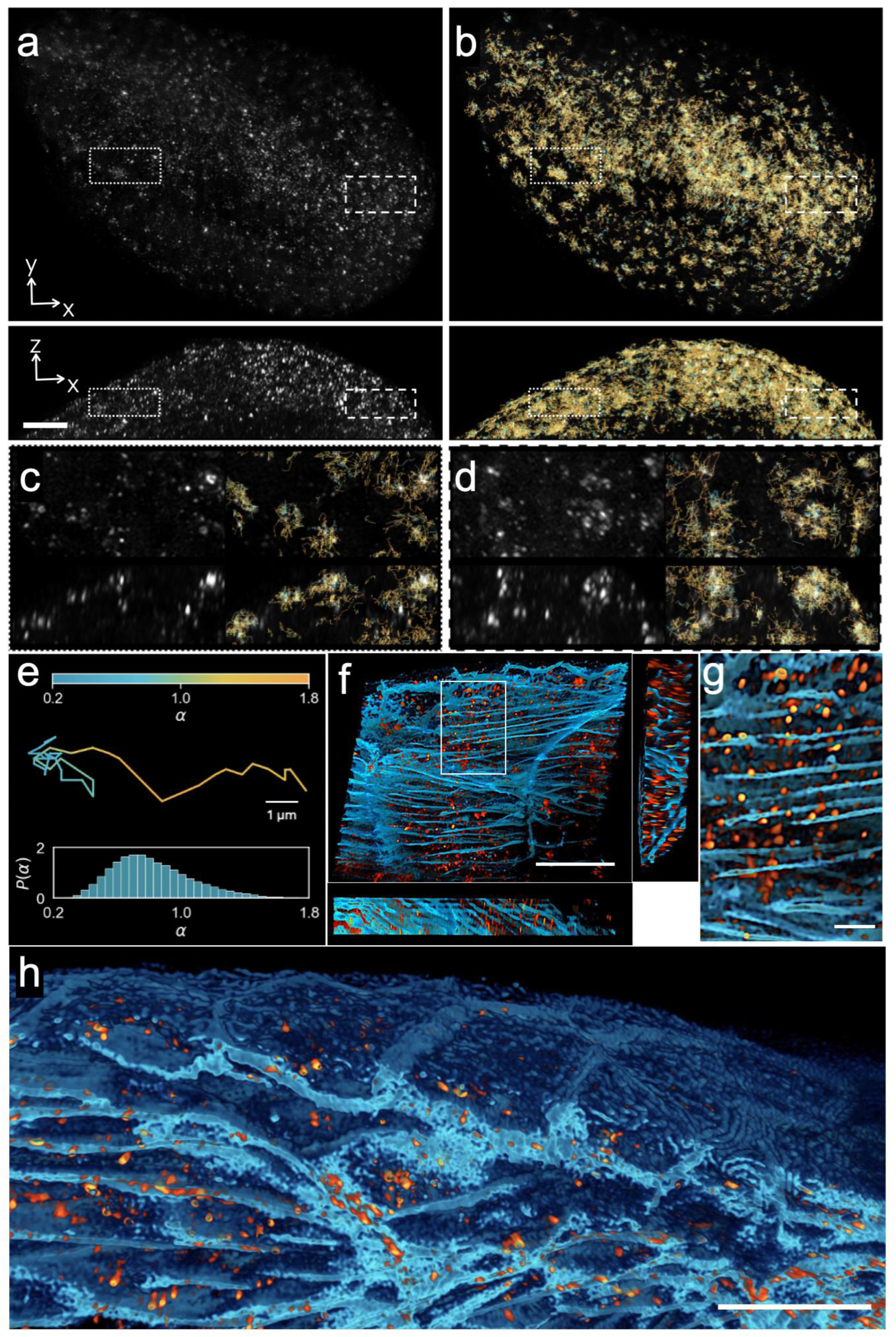
Endosomal motility in zebrafish embryo tail tissue. (a–d) Data were collected by imaging the tailbud region of a 14-somite stage from a zebrafish embryo expressing mKate2-rab5ab after mRNA injection. mKate2-rab5ab-labelled endosomes were detected in a volume spanning approximately 266 × 266 × 60 µm^3^ and imaged over 12 min at 5.9 s per volume. Max intensity projections taken from (*top*) XY plane and (*bottom*) YZ plane are shown. **(b)** Overlays of endosome tracks onto raw images as in **a**, showing all endosomes tracked for at least 10 frames (60 s). Track segments are coloured according to the anomalous diffusion exponent (α) calculated by fitting the mean squared displacement (MSD) for each track segment. **(c–d)** Selected volumes corresponding to dotted (**c**) and dashed (**d**) insets in **a** and **b**. Panels are as described in **a**. Each volume spans 20 × 40 × 15 µm^3^. **(e)** Distribution of observed values of α, illustrated by (*top*) a representative trajectory that exhibits a range of motion from directed to confined, coloured according to α, and (*bottom*) the probability density of α calculated for each track segment in **a**. **(f)** Volume projection of data collected by imaging the lateral tail region of a zebrafish embryo 32 hours post fertilisation expressing mKate2-rab5ab (orange) and membrane-mNeonGreen (cyan) after mRNA injection. **(g)** Zoom of the region highlighted in **f**. **(h)** 3D view of the tail region of the zebrafish embryo expressing mKate2-rab5ab (orange) and membrane-mNeonGreen (cyan), displaying ridges and complex cellular morphologies. Scale bars: **(a, b):** 25 μm, **(f):** 50 μm, **(g, h):** 10 μm.

### Cross-scale mapping of macromolecular myosin assemblies in *Drosophila* development

A classical model organism in developmental biology is *Drosophila*. A key step in *Drosophila* embryogenesis is cellularisation, which involves the conversion of a single-celled syncytium to a multicellular embryo. Following cellularisation of the embryo, zygotic transcription ensues and distinct cellular behaviours are observable. Within 5 minutes, through the mid-blastula transition, the cephalic furrow and the ventral furrow form, as well as the cellular blastoderm, which expresses a mitotic pattern controlled by *string* expression that begins at the precephalic region. Gene expression patterns orchestrate cellular force generation based on actomyosin contractility that drives epithelial sculpting. In parallel, cell divisions also contribute to elongation and macroscopic behaviours of the embryo tissue. The divisions begin at the procephalic region and proceed in a successive manner across the embryo (*18*). *Drosophila* non-muscle myosin II regulatory light chain (encoded by the *spaghetti squash* gene*, sqh*) plays a key role in cellularisation, furrow ingression, and basal closure; it localises to the cleavage furrows during anaphase in dividing cells and is required for cytokinesis (*19*). Following sqh-3x-GFP in *Drosophila* embryo allows mapping both divisions, as it localises to cytokinetic rings, as well as contractile machinery assemblies that drive tissue -wide movements and folds. We imaged *Drosophila* embryos expressing sqh-3x-GFP from the stage of syncytial blastoderm through to gastrulation. To demonstrate how our high-resolution imaging approach enables rapid biological processes to be observed across a large area, we focussed on the posterior end of the embryo. In this region, dramatic tissue elongation takes place while simultaneously traversing the posterior fold over itself. This region has been difficult to capture in 3D at high resolution owing to the fast nature of movements, with the first phase of elongation lasting ∼25 minutes.

As the sqh-3x-GFP signal disappears post-cellularisation, it reappears at cell-cell junctions before the onset of posterior midgut invagination. By imaging sqh-3x-GFP beginning with the movement of primordial germ cells through the posterior midgut (PMG) invagination for 60 min at one volume every 25 s, we could follow changes in cell boundaries, contractile assemblies, and cytokinetic rings (Fig. 5a–c, Supplementary movies 7,8). The ensemble directions of cell divisions could be mapped by following the orientation of the division rings (Fig. 5d, Supplementary fig. 6). Visualisation of the orientation of the cell divisions revealed complex patterns: at a depth of 27 μm from the ventral surface, we find divisions that are oriented along the anterior-posterior axis (Fig. 5f), while at 35 μm, cell divisions were symmetrically placed to the midline and oriented along the dorsal-ventral axis (Fig. 5g, Supplementary movie 9). It is interesting to note that the wave of divisions occurs soon after the rapid phase of germ band elongation (GBE) and are programmed to take place in a specific temporal sequence that is constant between embryos ( *18*). Consistent with this, we also observe patterns of divisions occurring on the posterior-ventral side of the embryo, as the rapid phase of GBE concludes (Supplementary movie 10). Furthermore, with just two imaging volumes, we captured the entire *Drosophila* embryo (Supplementary movie 11) at high spatial resolution and with a time resolution of 66 seconds. Therefore, simultaneously occurring events that occur across the entire embryo can be mapped. For example, as precellularisation concludes, sqh-3x-GFP signals immediately localise to the future ventral furrow and execute furrow formation; at the same time, the cells in the precephalic region divide, as visualised by the formation of cytokinetic rings (Supplementary fig. 7). Waves of division in the precephalic region, which occur concurrently with formation of cell boundaries after ventral furrow formation, can also be mapped (Supplementary movie 12). By imaging an entire Drosophila embryo, typically 400–500 μm in length and 150–200 μm in width, we can follow sqh-3x-GFP through cellularisation, the appearance of the ventral and cephalic furrows, germline extension, and the formation of segments, allowing us to capture the evolution of molecular assemblies of sqh-3x-GFP that execute force generation (macroscopic, intercellular) and cell division (intracellular, localising in the cytokinetic ring).

**Figure 5.**
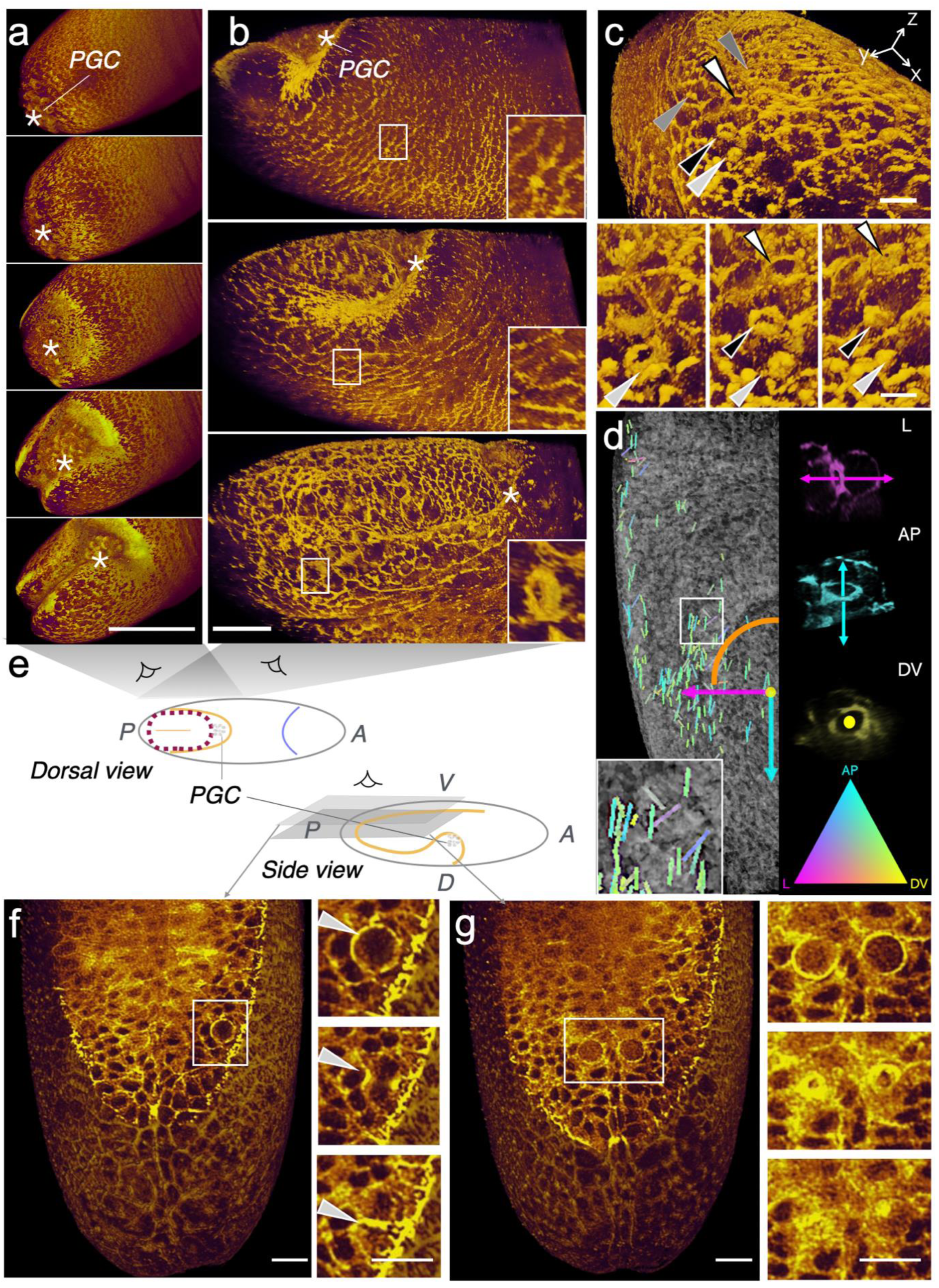
Correlation of rapid cellular dynamics and tissue -wide rearrangements in a *Drosophila* embryo expressing sqh-3x-GFP. **(a)** Timelapse imaging of posterior midgut (PMG) invagination shown every 1 min after initiation. (*) indicates primordial germ cells (PGC) **(b)** Side view of the PMG invagination at the posterior end of the embryo shown every 3 min. Cells appear to turn and elongate with the progress of invagination, subsequently undergoing cell divisions, inferred by formation of cytokinetic rings. **(c)** Divisions are also observed on the dorsal side of the extending tissue; arrows in the lower panel follow divisions (closing cytokinetic furrows). **(d)** *Left*: Transparent surface rendering of a *Drosophila* embryo during midgut invagination (orange arc) overlaid with division ring orientation vectors. Vectors are colour-coded by alignment with the lateral (L, magenta), anterior–posterior (AP, cyan), and dorsal–ventral (DV, yellow) axes. The white box indicates the magnified inset. *Right*: Representative division rings for each axis and the corresponding 3D orientation colormap. **(e)** Schematic describing the views presented in the images. **(f)** Cross-section at a depth of 27 µm from the ventral surface, displaying a cell division (in rectangle) oriented in the anterior-posterior axis. **(g)** Cross-section at a depth of 35 µm highlighting two cell divisions occurring symmetrically. In this case, the orientation is dorsal - ventral. Scale bars: **(a):** 100 μm, **(b):** 25 μm, **(c):** 10 μm, **(f, g):** 20 μm.

### Organelle dynamics in photosensitive mouse oocytes

To demonstrate our approach on a challenging highly photosensitive sample, we imaged mitochondrial streaming in mouse oocytes. In mouse oocytes, maturation is concomitant with cytoplasmic reorganisation. The meiosis II (MII) stage oocytes show a distinct accumulation of mitochondria in the spindle hemisphere, which displays a characteristic streaming ( *20–22*) in which the mitochondria move towards the spindle from the centre of the oocyte. Upon reaching the spindle, they move away from it, along the cortex. This flow pattern is halted at the equator, distinguishing the spindle hemisphere from the non-spindle hemisphere. The flow patterns occur throughout the 3D volume of the oocyte and are difficult to capture since oocytes are extremely sensitive to the light used for excitation, which can inhibit the streaming phenomenon. Therefore, only a few planes had previously been imaged to capture the prominent parts of the streaming patterns. Here, due to the photo-gentle nature of the light-sheet geometry, combined with the long non-diffractive length of the Airy beam that can traverse the diameter of the oocytes (typically 80 μm), we were able to capture the movements of mitochondria using mito-Dendra2 (*23*) throughout the 3D volume of the oocyte for up to 4 hours at a temporal resolution of 5 minutes (Fig. 6a, Supplementary movie 13). This enabled mapping of complex patterns of mitochondrial streaming within the oocytes (Supplementary fig. 8). This is illustrated by streamlines corresponding to 3D flow at distinct points in time, with sustained flows reaching speeds of ∼0.5 μm/min within the first 2 hours, which slow down then lose cross-oocyte coherence by 4 hours (Fig. 6b).

**Figure 6.**
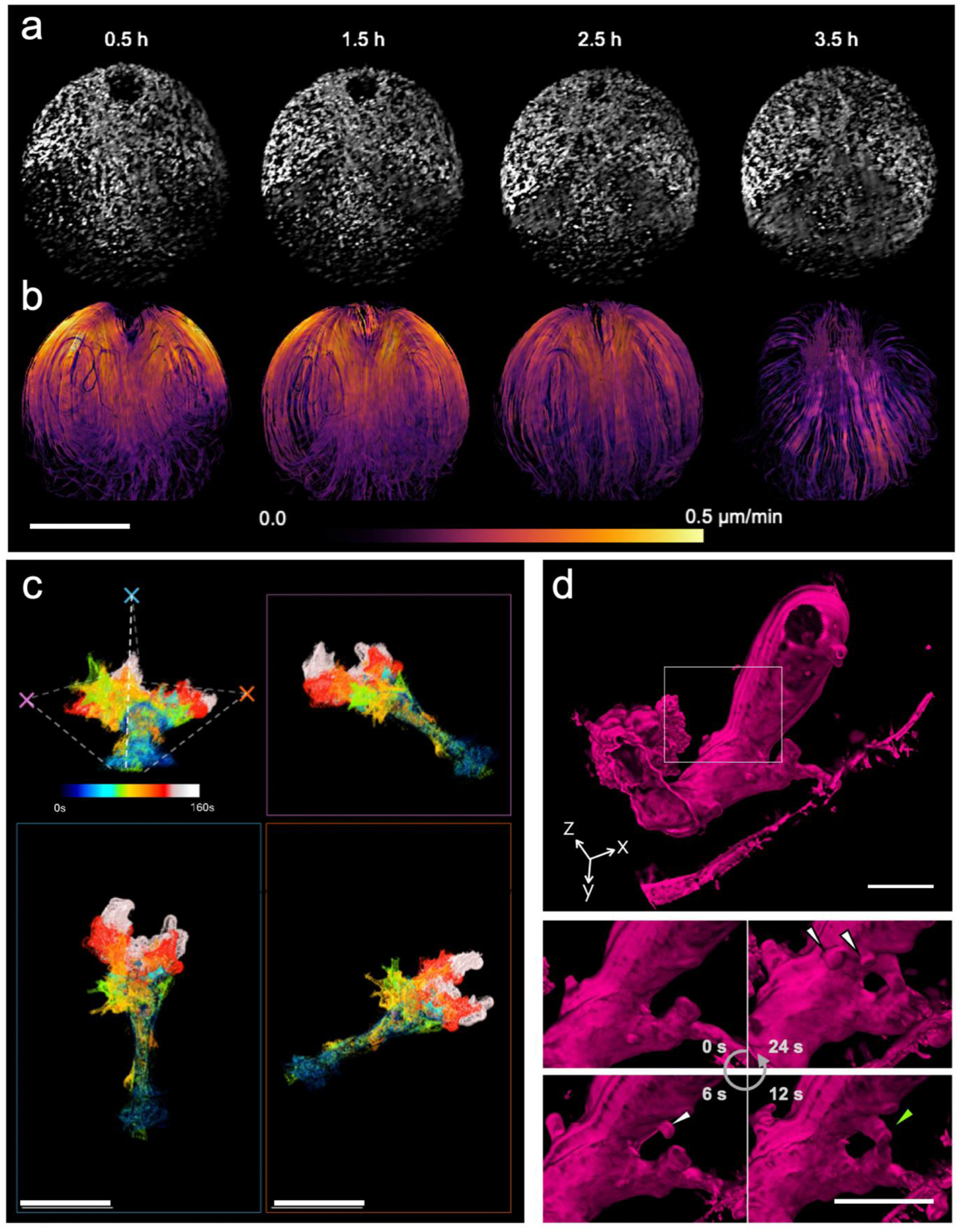
Quantification of cellular dynamics in large, photosensitive single cells. **(a)** Mouse meiosis II oocyte with labelled mitochondria (Dendra2) with diameter ∼80 μm was imaged every 5 min for 4 h. Max-intensity projections through the oocyte show the distribution of mitochondria at four equally spaced time points. **(b)** Maps of mitochondrial motion were calculated from optical flow. 3D flow maps of mitochondria throughout the entire oocyte volume were time-averaged over 60 min centred at the indicated timepoint. Time-average velocity at each voxel within the oocyte is colour-coded as indicated. **(c)** Multi-view temporal projections of an amoeba captured at 6.2 s per volume. Morphology changes over 160 s are colour-coded according to the temporal colour bar. The top-left orientation guide indicates the viewing angles (coloured crosses) corresponding to the perspectives shown in the magenta, cyan, and orange boxed panels. **(d)** Volumetric rendering of a segmented amoeba captured at 5.8 s per volume. The white box indicates the region shown in the time-lapse sequence below. White arrowheads mark extending protrusions and the green arrowhead highlights a fusion event, over a 24 -s interval. Scale bars: **(a,b)**: 50 μm, **(c):** 100 μm, **(d):** 50 μm.

### Fast, large-scale motility of amoebae

To capture another example of very large, dynamic single cells, we imaged *Amoeba proteus. A. proteus* cells form wide, thick pseudopods, referred to as lobopods, that enable amoeboid movement. These protrusions are thought to be driven by the creation of intracellular pressure arising from contractile actomyosin systems; at the site of elongation, the actin cytoskeletal structure collapses, causing cytoplasm to flow into the weakened path (*24*). A single *A. proteus* cell typically extends 250–750 μm, undergoes constant changes in cell shape, and exhibits an average crawling speed of 4 μm/s (*25*). Owing to its extremely large size and rapid dynamics, capturing the entire cell body with a single-volume acquisition is challenging. Taking advantage of the Airy beam’s ability to span the entire FoV (332 μm by 332 μm for a 20× 1.0 NA detection objective) and using large step sizes (1 μm between XZ planes), we were able to image a volume enclosing the entire cell body in 6.2 s (Fig. 6c, Supplementary fig. 9, Supplementary movie 14) and thereby capture the rapid protrusion formation and dynamics. We also observed two lobopodial extensions that fused at their distal ends (Fig. 6d, green arrow), resulting in a closed-loop morphology and a lumen within the surface of the amoeba.

### Organelle dynamics in cancer organoids

To demonstrate live imaging in multicellular assemblies, we imaged patient-derived colorectal (CRC) organoids embedded in Matrigel (*26*) (Fig. 7a,b). A longer Airy beam could provide subcellular resolution on single-cell thick hollow organoids but was limited in penetrating solid, non-hollow organoids as the beam quality was compromised by scattering. In the context of cancer, mitochondria are involved in multiple cellular processes that regulate tumour development, including metabolic reprogramming and metastasis. Mitochondrial localisation and morphologies are associated with metastatic potential (*27*). We captured mitochondrial dynamics including fusion and fission in individual cells, highlighting the ability to observe contemporaneous organelle dynamics across any sub-region within the organoid (Fig. 7c). Combined with screening approaches, using such volumetric imaging to monitor organelle dynamics has the potential to identify patterns that result in spontaneous metastasis.

**Figure 7.**
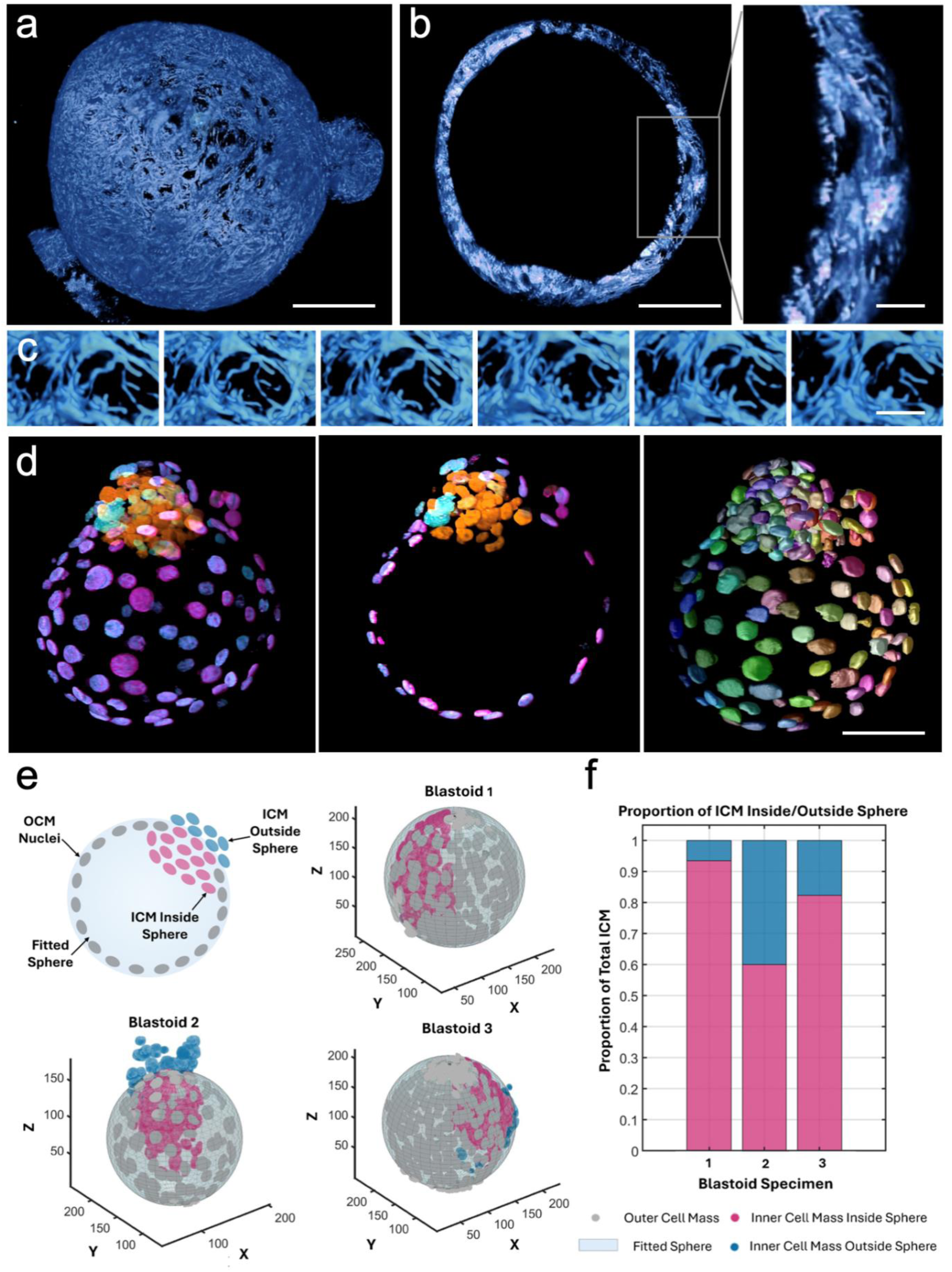
**(a)** PKmitoDeepRed-labelled patient-derived colorectal cancer organoid rendered in NVIDIA IndeX **(b)** Cross-section at the diameter of the cancer organoid showing single-cell-thick wall of the organoid. **(c)** Time-lapse montage of mitochondrial dynamics acquired every 64.6 seconds. **(d)** 3D raw rendering *(left)* and cross-sectional view *(middle)* of a representative iBlastoid stained for GATA3 (magenta), Nanog (orange), and GATA6 (cyan). *Right:* Corresponding instance segmentation of individual nuclei. **(e)** Cross-sectional schematic of iBlastoid analysis *(top left):* a sphere is fitted to the outer cell mass (OCM) nuclei (grey) to classify inner cell mass (ICM) nuclei as inside (pink) or outside (blue) the boundary. The remaining panels display representative 3D colour-coded segmentations of three blastoids analysed using this method. Coordinate units are in µm. **(f)** Quantification of the proportion of ICM nuclei located inside (pink) versus outside (blue) the fitted sphere for the three representative blastoids shown in **e**. Scale bars: **(a,b):** 50 μm, zoom: 10 μm; **(c):** 10 μm; **(d):** 50 μm.

### Rapid screening of iBlastoids

Blastocysts, the multicellular structures that develop into early embryos, can be modelled *in vitro* using iBlastoids (*28*). Spatial cell type profiling and localisation to extract patterns are necessary steps to characterise iBlastoids, typically accomplished by immunostaining analysis using markers for distinct cell types including GATA3 (trophectoderm), Nanog (epiblast), and GATA6 (primitive endoderm). iBlastoids are typically 100 μm in diameter and require imaging at sufficient sectioning and resolution to capture all cells, with multiple channels to distinguish individual cell type markers, the nucleus, and cell boundaries. Previously, confocal microscopy was used to acquire such volumes, which is time-consuming and thus prohibitive of high-throughput screening. Here, we could capture each iBlastoid in five different channels in ∼10 min, at a step size of 200 nm between each slice. This enabled us to visualise the detailed distribution of transcription factors (Fig. 7d, Supplementary movie 15) in iBlastoids and to use a morphological feature—the presence of inner cell mass outside a fitted sphere of the iBlastoid—as a parameter for quality assessment (Fig. 7e, f).

## Discussion

Light-sheet fluorescence microscopy enables imaging of cells and tissues across a wide range of length scales. However, current methodologies require a trade-off between volume of imaging, spatial resolution, and temporal resolution, where only two of any of these factors may be optimised within a single acquisition. Thus, a key challenge is to achieve simultaneous imaging from near-diffraction-limited structures up to tissue-level features, whilst retaining sufficient temporal resolution to capture subcellular dynamics. Here, we report the first use of Airy beam light-sheet microscopy in a high-NA configuration to capture large volumes of biological samples without significant loss of spatial or temporal resolution. To demonstrate the broad utility of this approach, we present applications across a range of systems and processes.

First, the ability to capture rapid dynamics at high resolution over a large volume is beneficial for the study of biological processes coordinated across scales, such as animal development. We illustrate simultaneous capture of organelle dynamics from large tissue samples from zebrafish embryos and larvae, as well as coordination of cellular division events with rapid, large -scale changes in tissue morphology in *Drosophila* embryos. Second, large 3D samples that require exceptionally photo-gentle imaging, such as oocytes, benefit from the ability to capture complete volumes without the need for tiling. Third, this technique permits the capture of extremely rapid events occurring at or above the cellular scale, as illustrated by movies of pressure-driven morphological changes or stochastic lobopodial extensions across the entire surface of an amoeba. Finally, this approach is suitable for high-throughput screening of large, multicolour samples including cell lines, cancer organoids, and iBlastoids, which is often time-prohibitive using standard methodologies.

One limitation of Airy beam imaging is the highly asymmetric PSF, which necessitates that all data be deconvolved prior to visualisation and analysis. We offset this disadvantage through the design of an XZ-scan that does not require the additional deskew step typical for similar setups. We also perform deconvolution with PSFs obtained for specific beams on a high-performance computing cluster to minimise the resultant time delay. Another limitation is that single photon excitation restricts imaging in deeper parts of samples that are highly scattering, such as solid cancer organoids. This can be overcome using multiphoton excitation, which has been demonstrated with Airy beam light-sheet microscopes (*29, 30*), albeit with low-NA objective systems. In addition, while the use of Airy beams offers extended FoVs, it does limit depth-dependent correction using adaptive optics as has been demonstrated for lattice light-sheet microscopy (*11*). However, using shorter Airy beams provides the versatility to implement similar approaches. The most significant post-acquisition challenge lies in the size of the datasets generated, which can reach >1 TB for a single volume. As such, adequate computational infrastructure must be in place to transfer, store, visualise, and analyse these very large multidimensional datasets. New approaches such as hierarchical image visualisation are needed to efficiently load images and allow users to interact in 3D with datasets where a single volume can easily exceed available memory. PetaKit5D, for example, is a tool that mitigates many such issues associated with large datasets ( *31*). This is required both to qualitatively interpret the phenomena captured, as well as to guide rigorous quantitative analysis.

The ability to image complex biology across a wide range of spatial and temporal scales provides a significant advancement toward deciphering emergent phenomena that span multiple levels of organisation. Instead of building a description of dynamic events from datasets acquired using distinct modalities for different samples under varying conditions, Airy beam light -sheet microscopy permits simultaneous acquisition of processes ranging from organelle dynamics to tissue-level rearrangements within a single live biological specimen. This opens new opportunities to link mechanisms operating at the molecular and subcellular levels to concomitant higher-level processes, such as active growth and patterning in developing tissues. Large imaging volumes also directly entail acquisition of large numbers of individual events operating at small, rapid scales, such as protein movements and organelle interactions, which is needed to establish mechanism in highly stochastic processes. This approach carries significant potential, especially in the field of developmental biology, to accelerate efforts to understand how biochemical events drive robust processes such as cell differentiation and tissue patterning (*32–34*).

## Materials and Methods

### Optics Configuration

Light-sheet imaging was performed using a custom-designed Aurora Airy Beam Light-Sheet Imaging System (build outsourced to M-Squared). This upright light-sheet microscope utilises an asymmetric orthogonal high-NA objective configuration with a 20× 0.6 NA (Thorlabs) excitation objective and a 25× 1.1 NA (Nikon) or 20× 1.0 NA (Olympus) detection objective. Six laser lines with wavelengths in the visible spectrum (405, 445, 488, 561, 647 and 685 nm, respectively) are available for excitation. The Airy beam is generated by applying a cubic phase mask on the collimated beam (9 mm diameter) using a reflective spatial light modulator (SLM, Meadowlark MSP1920 400-800). The SLM modulates the wave front at the back aperture with a cubic polynomial function: 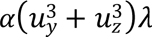, where *u*_*y*_ and *u_z_* are the normalised Cartesian pupil coordinates aligned with the *y*- and *z*-axes respectively, and λ is the excitation wavelength. The dimensionless parameter α dictates the propagation invariance of the Airy beam. The beam is demagnified by a factor of 0.375, and the conjugate SLM plane is projected onto the galvanometer mirror. The beam is scanned using the mirror corresponding to the *x*-direction, located after the excitation objective, to create a scanned light sheet. The beam is then magnified by a factor of 1.125 compared to the SLM. After Fourier transform, the propagation length is measured with respect to the camera field of view (FoV) to define the Airy beams. For example, Airy 5 corresponds to 5% of 266 µm, which is the length of the side of the camera FoV for a 25× 1.1 NA objective. A second galvanometer mirror is used to position the beam in the plane of the focus of the collection objective. In the detection path a dichroic mirror is used to split emission light of different wavelengths to two sCMOS cameras (Hamamatsu Orca Flash 4.0 V3) enabling sequential multi- or simultaneous dual-channel acquisitions. All parameters of imaging of various samples are summarised in Supplementary table T2.

### Zebrafish Husbandry

Zebrafish (*Danio rerio*) husbandry and breeding was conducted in the AquaCore facility at Monash University according to standard procedures (*34*). All experimental procedures were approved by the Monash University Animal Ethics Committee under ethics approval numbers ERM22161 and ERM41803. Embryos were reared at 28 °C or 22 °C in E3 media (5 mM NaCl, 0.17 mM KCl, 0.33 mM CaCl_2_, 0.33 mM MgSO_4_) until they reached the appropriate developmental stage. Larvae older than 24 hours post fertilisation (hpf) were treated with 75 µM PTU (1 -phenyl-2-thiourea) to prevent pigmentation. Ethyl-*m*-aminobenzoate methanesulfonate (Tricaine) was used to anaesthetise embryos and larvae (0.168 mg/mL) and adult fish (0.3 mg/mL), where necessary. Zebrafish wild type strains Tuebingen (TU) and AB as well as the transgenic line *Tg(actc1b:mCherry-CAAX)^pc22Tg^* (*35*) were used in this project. To maintain genetic diversity, transgenic lines were outcrossed to wild type strains every second generation.

### Generation of plasmids and transgenic strains

Transgenic constructs were assembled using the multisite gateway cloning kit ( *36*). The muscle-specific mito-GFP construct (*actc1b*:mito-GFP) was generated as per (*37*). The muscle-specific ER-mCherry construct (*actc1b*:ER-mCherry) was generated using p5E-*actc1b* (*36, 37*), pME-mCherry-ER-3, which was subcloned from a plasmid gifted by Michael Davidson (Addgene plasmid # 55041; http://n2t.net/addgene:55041; RRID:Addgene_55041), p3E-pA and pDEST-Tol2-pA2 (*36*). Plasmids were injected at 30 ng/μL into 1-cell-stage embryos along with transposase RNA (25 ng/μL) that was synthesized from the pcs2FA-transposase vector using the mMessage machine Sp6 kit (Ambion, AM1340). The final transgenic lines created were: *Tg(actc1b:mito-GFP)^uom407Tg^* and *Tg(actc1b:ER-mCherry)^uom408Tg^*.

### mRNA Microinjections

Prior to injection, plates containing 3% agarose imprinted with grooves were prepared by placing a Tu-1 microinjection mould into liquid agarose. Microinjection needles were prepared using a P-2000 micropipette puller to pull needles from glass capillaries (1 mm outer diameter, 0.78 mm inner diameter). A standard microinjection apparatus was used to inject 1 nL of 50 –100 ng/µL capped mRNA in nuclease free water, supplemented with 10% phenol red as injection guide, into zebrafish embryos at the single-cell stage. The relevant mRNA was prepared from linearised plasmid DNA containing a SP6 promoter site by in-vitro transcription using the mMessage mMachine SP6 Transcription Kit (Invitrogen). The plasmid for Membrane-mNeonGreen was a gift from Amro Hamdoun (Addgene plasmid # 198057; http://n2t.net/addgene:198057; RRID:Addgene_198057). The PC2+-mKate2-rab5ab was generated by GenScript by cloning the mKate2-rab5ab sequence from a reference plasmid into the PC2+ backbone. The actc1b-mKate2-rab5ab reference plasmid was a gift from Rob Parton (Addgene plasmid # 109649; http://n2t.net/addgene:109649; RRID:Addgene_109649).

### Zebrafish Embryo and Larva Preparation for Live Imaging

Before mounting, zebrafish embryos/larvae were manually dechorionated and anaesthetised using 168 mg/L Tricaine in methylene blue-free E3, where necessary. All further steps were carried out using methylene blue-free E3 media.

The embryos and larvae were mounted in a volcano-shaped mount placed inside the microscope’s imaging dish. To produce a volcano-shaped mount, 1.2% low melting point agarose in E3 was shaped using a mould (based on (*11*)). Live anaesthetised embryos/larvae were transferred to melted 0.8% low melting point agarose in E3 at 42 °C and then transferred into the volcano mount, adjusting their orientation before the agarose solidified. After the agarose solidified, the imaging dish was filled with E3 media containing 168 mg/L Tricaine for imaging.

### Drosophila

*D. melanogaster* were raised at room temperature (22–23 °C) or 18 °C on food made with yeast, glucose, agar and polenta. Animals were fed in excess food availability to ensure that nutritional availability was not limiting. All experiments were carried out at 25 °C. Males and females were used for all experiments. Mounting was performed on 5 mm coverslips coated with glue extract prepared by leaving double-side sticky tape in hexane overnight. The following genotype was used: *sqh-3x-GFP*.

*Drosophila* crosses were established at 25°C on apple juice plates with yeast paste in embryo collection cages. Embryos of the appropriate stage were collected from apple juice plates using a wet paintbrush, washed in an embryo strainer with deionised water and manually dechorionated using Dumont no. 5 forceps on double sided tape. Dechorionated embryos were adhered to the imaging sample holder using folded double-sided tape to create an angled surface, allowing appropriate orientation of embryos for image acquisition. Embryos were submerged in deionised water for the duration of the imaging session.

### Mouse Oocytes

All animal experiments in this study were approved by the Monash University Animal Ethics Committee and conducted in accordance with the Australian National Health and Medical Research Council (NHMRC) Guidelines on Ethics in Animal Experimentation.

7-week-old PhAM (photo-activatable mitochondria) mice (*23*) were superovulated by intraperitoneal injection of 5 IU of pregnant mare’s serum gonadotropin (Prospec) followed 44 – 48 h later by intraperitoneal injection of 5 IU of human chorionic gonadotropin (hCG) (MSD Animal Health). 12–13 h after hCG injection, oviductal cumulus masses were released into pre-warmed M2 medium (Sigma-Aldrich) supplemented with 300 µg/mL hyaluronidase (Sigma-Aldrich) to remove cumulus cells. Oocytes displaying a first polar body (indicating metaphase II arrest) were washed and transferred to drops of M2 medium under mineral oil.

These oocytes were microinjected with mRNA using an electrophysiology-based picopump (PV820, World Precision Instruments) and a micromanipulator (MMN-1, Narishige). Following microinjection, oocytes were incubated in M2 medium under mineral oil (RT, 10 min) before being transferred to a heat block (37 °C, 10 min) to facilitate oocyte retrieval. Oocytes were cultured in M2 medium for at least 3 h for transgene expression.

### Amoeba Sample Preparation and Mounting

Amoebae were cultured on 5 mm glass coverslips submerged in protist culture medium (Southern Biological) supplemented with five grains of rice. The cultures were maintained in the dark at room temperature for 5–7 days prior to imaging. For live imaging, coverslips were mounted onto a raised platform sample holder inside the imaging dish, which was filled with protist culture medium containing the lipophilic dye Fast DiI Solid (Thermo Fisher) at a final concentration of 2.5 µM. Imaging commenced 10 min post-staining without subsequent media replacement.

### Cancer organoids

Patient-derived colorectal cancer (CRC) organoids were established as previously ( *26*) described and in accordance with the Declaration of Helsinki, and the protocol was approved by the Cabrini Research Governance Office (CRGO04-19-01-15) and the Monash Human Research Ethics Committee (MHREC ID 2518). TUBB::TagGFP2 CRC organoids lines were then generated using CRISPR-HOT(*39*) with CRISPaint gene tagging Kit (Addgene #1000000086) (*38*) and an sgRNA (5’-gaggccgaagaggaggccta-3’) plasmid (Genscript). 10 days post-passage organoids were mixed at a 1:1 (v/v) ratio with TrypLE-passaged organoids and resuspended in Matrigel (Corning) containing 1:50,000 fluorescent beads. A 10 µL droplet of the suspension was seeded onto a UV-sterile Parafilm on ice and immediately covered with a 5 mm round coverslip. After the Matrigel solidified for 10 min in 37 °C, the coverslip with the Matrigel disc was carefully lifted from the Parafilm and placed into a 6-well tissue culture plate (Nunc) with the Matrigel layer facing upward. 2 mL of phenol-red-reduced complete CRC organoids culture medium (*26*) supplemented with 10 μM Y-27632 dihydrochloride kinase inhibitor (Tocris Bioscience) were added to each well. The following day, coverslips were incubated with PKmitoDeepRed 10 nM for 20 min. For imaging the coverslip was mounted on a raised platform sample holder inside the imaging dish, which was filled with pre-warmed culture media.

### iBlastoids

iBlastoids were generated according to established protocols (*28*). iBlastoids were collected into a protein low binding tube under a dissecting microscope. After washing once with PBS by centrifuging for 1 min at 10–20 × g, iBlastoids were fixed in 4% PFA for 40 min, washed with PBS and permeabilised with 0.1% Triton X-100 (Sigma) in PBS for 20 min, then blocked with 10% donkey serum (Thermo Fisher). Primary antibodies used were rabbit anti-NANOG polyclonal (1:100, Abcam, ab21624), mouse anti-GATA3 (1:100, BD Biosciences, 558686) and goat anti-GATA6 (R&D AF1700). Primary antibody incubation was conducted overnight at 4 °C on shakers followed by incubation with secondary antibodies (donkey anti rabbit 488, donkey anti mouse 555, donkey anti goat 647, 1:500, Thermo Fisher). After labelling, iBlastoids were stained with 4*′*,6-diamidino-2-phenylindole, dihydrochloride (DAPI) (1:1000, Thermo Fisher) for 10 min. iBlastoids were stained with phalloidin (A22286, Thermo Fisher) for 1 h before imaging. Stained iBlastoids were transferred into an FEP tube (FT0.8X1.0 FEP UTW, Adtech). Tubes were sealed with grease and mounted in the imaging chamber, which was filled with PBS.

### Cell Culture and Mounting

HeLa Rab5-GFP Rab7-mCherry and RPE1 ER-StayGold HaLo-Mito cells were incubated at 37 °C in 5% CO_2_ in high glucose Dulbecco’s modified Eagle’s medium (DMEM) (Life Technologies), supplemented with 10% foetal bovine serum (FBS) and 1% penicillin and streptomycin (Life Technologies). Cells were seeded at a density of 200,000 cells per well in a six-well plate containing 5 mm glass coverslips. RPE1 ER-StayGold HaLo-Mito cells were incubated in 50nM JFX650 for 30 min, followed by a PBS wash and media replacement prior to imaging. For imaging the coverslip was mounted on a raised platform sample holder inside the imaging dish. All cells were imaged in phenol red-free DMEM heated to 37 °C.

### Resolution Maps

Resolution maps were calculated for each plane of an image volume using single image Fourier ring correlation (*15*). Images corresponding to single cells consisted of a thin strip near the glass coverslip, scanned across the width of the FoV (using an acquisition mode where each variation in depth corresponds to lateral movement through the sample). The location of glass–cell interface was identified by segmentation, and at each image depth, a resolution map was generated by calculating the resolution within subregions of 256 × 256 pixels using a rolling window over a column spanning 256 pixels in width that covers the cells without incorporating blank regions far from the glass interface. The full resolution map was collected by aggregating each local value of the resolution at each depth (corresponding in this case to specific positions along the *X* axis within the final image volume). Resolution maps for zebrafish tissue were calculated by tiling subregions spanning 256 × 256 pixels (i.e., 64 discrete subregions at each depth). Resolution maps were calculated using the original 1FRC MATLAB implementation (*15*), supplemented by custom MATLAB code for tiling and rolling window analyses. For visualisation, final resolution maps were smoothed using a Gaussian filter and grayscale data were assigned RGB values corresponding to the local value of the resolution using custom Python code, prior to visualisation using napari ( *39*).

### Endosome Tracking and Analysis

Endosomes were detected and tracked as previously reported (*17*) using custom Python code. Briefly, a Laplacian of Gaussian filter was used to detect individual endosomes within each frame, and complete trajectories were constructed using Trackpy. To analyse endosomal motility, a rolling window mean-squared displacement (MSD) was conducted using custom Python code. Briefly, all trajectories were dedrifted, then the MSD was calculated for segments of each trajectory spanning at least 10 frames (60 s), and the anomalous diffusion exponent (α) extracted for all segments for which the lag time and MSD fit well to a line on a log–log plot. The resulting time-dependent value of α was then smoothed using a Savitzky-Golay filter.

### Oocyte Flow Maps

An image stack containing a single oocyte was first dedrifted by using phase cross-correlation to account for translational drift due to temperature fluctuations across the movie, as well as histogram equalisation to account for variations in image intensity over time. The optical flow was then calculated for each pair of subsequent frames using the iterative Lucas-Kanade (iLK) algorithm as implemented in scikit-image. To smooth out fluctuations, a time-average flow map was constructed by averaging four contiguous 3D flow maps (corresponding to 60 min of imaging), prior to calculating 3D streamlines for visualisation. All analyses were conducted using custom Python code.

### Amoeba Visualisation

Raw volumetric image data of DiI-stained amoebae underwent initial pre-processing using a custom semi-automatic pipeline in Fiji to remove coverslip signal. Segmentation was achieved by training a custom machine learning-based pixel classifier in Labkit. Downstream analysis employed custom scripts (MATLAB 2023b) for post-processing the segmentation output, specifically to isolate the largest object and thus exclude smaller, non-target microorganisms present in the culture. Volumetric rendering of the segmented amoebae was performed using the 3DScript plugin in Fiji. Finally, custom MATLAB scripts were used to temporally colour code the segmented amoeba volumes for visualisation. Final visualisations were generated by projecting the temporally coloured volumes onto raw data extracted regions.

### Zebrafish Myotome Analysis

Segmentation masks for myotomes and nuclei were generated in MATLAB utilizing the Medical Imaging Toolbox Interface for Cellpose Library support package (*40*). Prior to segmentation, the volumetric dataset of the endoplasmic reticulum (ER) channel was scaled down using a custom MATLAB code such that the final mean equivalent diameter of the object of interest was 50 pixels. Nuclei segmentation employed a custom Cellpose model trained using ten image slices, evenly distributed throughout the volume, that were manually annotated in Labkit. Nuclear boundaries were specifically defined by annotating the signal-devoid spaces within the myotomes. Segmentation of mitochondria and ER was achieved using a machine learning-based pixel classifier trained in Labkit for each channel respectively. The resulting confidence maps were subsequently post-processed via a custom MATLAB script to threshold the signal and remove false positives arising from noise, based on object size criteria. For ER and mitochondrial orientation analysis, the segmentation output of the tissue’s centre slice was skeletonised. Orientations of individual branches were computed using the Image Processing Toolbox in MATLAB, and the orientation distribution was displayed using representative semi-polar plots generated by custom-written MATLAB code.

### iBlastoid Nuclei Extraction and Analysis

iBlastoid volumes were first pre-processed with channel registration using Fiji plugin Fast4DReg to ensure precise signal localisation of all channels. Nuclei were segmented using a custom-trained model built upon the Cellpose-SAM model (*40*). The model training involved manual annotation on randomly generated XY, YZ, and XZ slices. Segmentation outputs were validated for accuracy by visualising overlays onto the raw data using syGlass with an Oculus VR headset. Inner and outer cell masses were manually annotated in VR. To quantify nuclei distribution of the inner cell mass, custom MATLAB code was written. Least squares fitting was utilised to generate the best-fit sphere which encapsulated the outer cell mass. The proportion of inner cell mass cells inside and outside of this fitted sphere was then quantified.

#### *Drosophila* Embryo Division Orientation Analysis

*Drosophila* embryo volumes were segmented using a machine learning-based pixel classifier trained in Labkit. Division rings at relevant timepoints were manually identified and annotated using syGlass and an Oculus VR headset. The resulting division ring masks were exported and processed using custom MATLAB code. The ring’s orientation was calculated by applying Principal Component Analysis (PCA) to the mask voxels where each division ring’s orientation was defined by the normal vector (*N*), corresponding to the third principal component. The anterior-posterior (AP) axis endpoints of the embryo were manually defined using VR, which was then used to establish an orthogonal 3D coordinate system relative to the embryonic axes: anterior-posterior (*V_AP_*), lateral (*V_L_*), and dorsal-ventral (*V_DV_*). *N* vectors extracted across several timepoints were mapped onto this coordinate system and colour-coded using a continuous Cyan, Magenta, Yellow (CMY) colormap. The final colour of a vector was derived from the normalized absolute dot product of the individual *N* vector with the *V_AP_*, *V_L_*, and *V_DV_* axes, respectively. This determined the weighting of the respective colours, with Cyan indicating complete AP alignment, Magenta indicating complete L alignment, and Yellow indicating complete DV alignment. A single representative timepoint was extracted and segmented using Labkit to finally generate a surface rendered visualization in MATLAB, which was overlaid onto the same 3D coordinate system as the orientation vectors.

### NVIDIA IndeX Visualisation

NVIDIA IndeX (https://developer.nvidia.com/index) was deployed to visualise movies of 0.5–3 TB in real time on 4× H100 Nvidia GPUs, Intel 72 Core CPU, and 1 TB RAM with 1 PB network storage.

## Acknowledgements

S.A. is supported by The EMBL Australia Partnership Laboratory (EMBL Australia) under the National Collaborative Research Infrastructure Strategy of the Australian Government. The authors thank Monash BDI Advanced Bioimaging. The Australian Regenerative Medicine Institute is supported by grants from the State Government of Victoria and the Australian Government. S.A. acknowledges NVIDIA for sharing NVIDIA IndeX to visualise our large datasets. The computational analysis and visualisation with NVIDIA IndeX were supported by Monash eResearch capabilities, including M3 High Performance Computing. S.A. acknowledges Wellcome Trust Team Science Grant. K.F.H. was supported by grants from the NHMRC (APP1194467) and ARC (DP230101406 and DP 250103072). A.A.R. was supported by a grant from ARC (DP240102721), P.D.C. was supported by grants from the NHMRC (GNT2016338) and ARC (DP240101647 and DP240102156). J.K. is funded by NHMRC Ideas grants GNT2037953, ARC Discovery Project Grant DP210103501, ARMI Accelerator Fellowship, Research Council of Finland, and the Sigrid Juselius Foundation and Biocenter Finland. I-W.L. and J.C. are funded by the ARC DP160104892 and NHMRC 1165627 and 200112. This work was additionally supported by NHMRC project grants APP1104560 and APP2004627 to J.M.P. The authors acknowledge Prof Paul McMurrick and the colorectal surgeons at the Cabrini Monash Department of Surgery for their contributions to specimen collection. S.A. would like to thank Srigokul Upadhyayula (University of California, Berkeley) for discussions and advice on data handling and visualisation.

## Author contributions

Project supervision: SA

Biological reagents, and sample preparation: CSW, SU, LZK, HRG, AP, HMY, SS, SAM,GS, I-WL, WHC, EB, SH, SC, HEA, JK, PC, KFH, JMP, JC

Analysis software: CSW, AP, SU, SA

Formal analysis: CSW, SU, SA

Data visualisation, figure and movie preparation: SA, AP, SU, CSW

Manuscript preparation: SA, CSW with input from all coauthors

## References

1. Y. Chen et al., Bridging single cells to organs: Mesoscale modules as fundamental units of tissue function. Cell 188, 6393–6410 (2025).

2. A. N. Kapanidis et al., From sequence to function: Bridging single-molecule kinetics and molecular diversity. Science 391, 458–465 (2026).

3. L. Z. Kreplin, S. Arumugam, High-resolution light-sheet microscopy for whole-cell sub-cellular dynamics. Current Opinion in Cell Biology 85, 102272 (2023).

4. W. C. Lemon, K. McDole, Live-cell imaging in the era of too many microscopes. Current Opinion in Cell Biology 66, 34–42 (2020).

5. P. J. Keller et al., Fast, high-contrast imaging of animal development with scanned light sheet-based structured-illumination microscopy. Nat Methods 7, 637–642 (2010).

6. M. Lange et al., A multimodal zebrafish developmental atlas reveals the state-transition dynamics of late-vertebrate pluripotent axial progenitors. Cell 187, 6742–6759.e6717 (2024).

7. P. Strnad et al., Inverted light-sheet microscope for imaging mouse pre-implantation development. Nat Methods 13, 139–142 (2016).

8. K. McDole et al., *In Toto* Imaging and Reconstruction of Post-Implantation Mouse Development at the Single-Cell Level. Cell 175, 859-876.e833 (2018).

9. C. Wolff et al., Multi-view light-sheet imaging and tracking with the MaMuT software reveals the cell lineage of a direct developing arthropod limb. Elife 7, (2018).

10. V. M. S. Ruetten et al., Imaging cellular activity simultaneously across all organs of a vertebrate reveals body-wide circuits. bioRxiv, 2025.2008.2020.670374 (2025).

11. T. L. Liu et al., Observing the cell in its native state: Imaging subcellular dynamics in multicellular organisms. Science 360, (2018).

12. T. A. Planchon et al., Rapid three-dimensional isotropic imaging of living cells using Bessel beam plane illumination. Nat Methods 8, 417–423 (2011).

13. B. C. Chen et al., Lattice light-sheet microscopy: imaging molecules to embryos at high spatiotemporal resolution. Science 346, 1257998 (2014).

14. T. Vettenburg et al., Light-sheet microscopy using an Airy beam. Nature Methods 11, 541–544 (2014).

15. B. Rieger, I. Droste, F. Gerritsma, T. Ten Brink, S. Stallinga, Single image Fourier ring correlation. Opt Express 32, 21767–21782 (2024).

16. D. Li, E. Betzig, Response to Comment on “Extended-resolution structured illumination imaging of endocytic and cytoskeletal dynamics”. Science 352, 527–527 (2016).

17. H. M. York et al., Deterministic early endosomal maturations emerge from a stochastic trigger-and-convert mechanism. Nature Communications 14, 4652 (2023).

18. V. E. Foe, Mitotic domains reveal early commitment of cells in Drosophila embryos. Development 107, 1–22 (1989).

19. R. E. Karess et al., The regulatory light chain of nonmuscle myosin is encoded by spaghetti-squash, a gene required for cytokinesis in Drosophila. Cell 65, 1177–1189 (1991).

20. K. Yi et al., Dynamic maintenance of asymmetric meiotic spindle position through Arp2/3-complex-driven cytoplasmic streaming in mouse oocytes. Nature Cell Biology 13, 1252–1258 (2011).

21. J. Van Blerkom, M. N. Runner, Mitochondrial reorganization during resumption of arrested meiosis in the mouse oocyte. Am J Anat 171, 335–355 (1984).

22. C. M. Dalton, J. Carroll, Biased inheritance of mitochondria during asymmetric cell division in the mouse oocyte. J Cell Sci 126, 2955–2964 (2013).

23. A. H. Pham, J. M. McCaffery, D. C. Chan, Mouse lines with photo-activatable mitochondria to study mitochondrial dynamics. Genesis 50, 833–843 (2012).

24. M. Dembo, Mechanics and control of the cytoskeleton in Amoeba proteus. Biophysical Journal 55, 1053–1080 (1989).

25. P. Soneji, E. J. Challita, S. Bhamla, Trackoscope: A low-cost, open, autonomous tracking microscope for long-term observations of microscale organisms. PLoS One 19, e0306700 (2024).

26. R. M. Engel et al., Patient-Derived Colorectal Cancer Organoids Upregulate Revival Stem Cell Marker Genes following Chemotherapeutic Treatment. J Clin Med 9, (2020).

27. L. Minarrieta et al., Mitochondrial elongation impairs breast cancer metastasis. Science Advances 10, eadm8212.

28. X. Liu et al., Modelling human blastocysts by reprogramming fibroblasts into iBlastoids. Nature 591, 627–632 (2021).

29. X.-J. Tan et al., Volumetric two-photon microscopy with a non-diffracting Airy beam. Opt. Lett. 44, 391–394 (2019).

30. N. A. Hosny et al., Planar Airy beam light-sheet for two-photon microscopy. Biomed Opt Express 11, 3927–3935 (2020).

31. X. Ruan et al., Image processing tools for petabyte-scale light sheet microscopy data. Nature Methods 21, 2342–2352 (2024).

32. M. Kirschner, J. Gerhart, T. Mitchison, Molecular “Vitalism”. Cell 100, 79–88 (2000).

33. A. M. Turing, The chemical basis of morphogenesis. Philosophical Transactions of the Royal Society of London. B, Biological Sciences 237, 37–72 (1952).

34. P. Liberali, A. F. Schier, The evolution of developmental biology through conceptual and technological revolutions. Cell 187, 3461–3495 (2024).

35. J. Berger et al., Loss of Tropomodulin4 in the zebrafish mutant trage causes cytoplasmic rod formation and muscle weakness reminiscent of nemaline myopathy. Dis Model Mech 7, 1407–1415 (2014).

36. K. M. Kwan et al., The Tol2kit: a multisite gateway-based construction kit for Tol2 transposon transgenesis constructs. Dev Dyn 236, 3088–3099 (2007).

37. A. A. Ruparelia, et al., Atrogin-1 promotes muscle homeostasis by regulating levels of endoplasmic reticulum chaperone BiP. JCI Insight 9, (2024).

38. J. L. Schmid-Burgk, K. Höning, T. S. Ebert, V. Hornung, CRISPaint allows modular base-specific gene tagging using a ligase-4-dependent mechanism. Nature Communications 7, 12338 (2016).

39. n. c. (2019), napari: a multi-dimensional image viewer for python.

40. C. Stringer, T. Wang, M. Michaelos, M. Pachitariu, Cellpose: a generalist algorithm for cellular segmentation. Nat Methods 18, 100–106 (2021).

