## supplementary figure for "Observing concurrent subcellular dynamics in large living tissues"

**[†]Equal first authors**

### Supplementary figure 1:

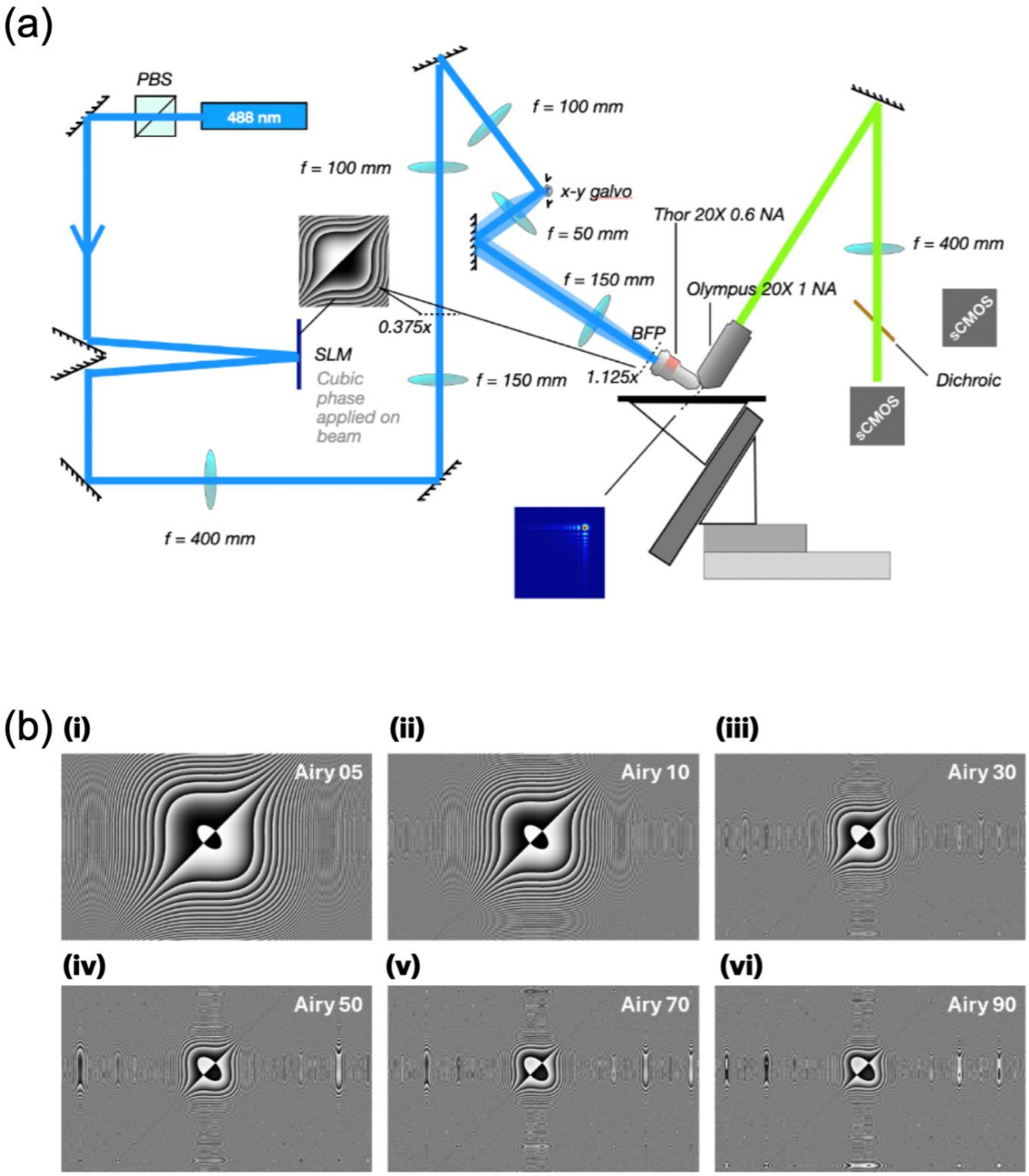


**Supplementary figure 1: Overview of the Airy beam light-sheet microscope**. **(a)** Optical layout of the Airy beam light sheet with the spatial light modulator (SLM) in the Fourier plane of the sample. Beam magnification factors at the SLM and the back focal plane (BFP) of the excitation objective are indicated. The resulting Airy beam profile along the axis of propagation is shown. **(b)** Patterns of phase masks used for various Airy beams in the manuscript.

### Supplementary figure 2:

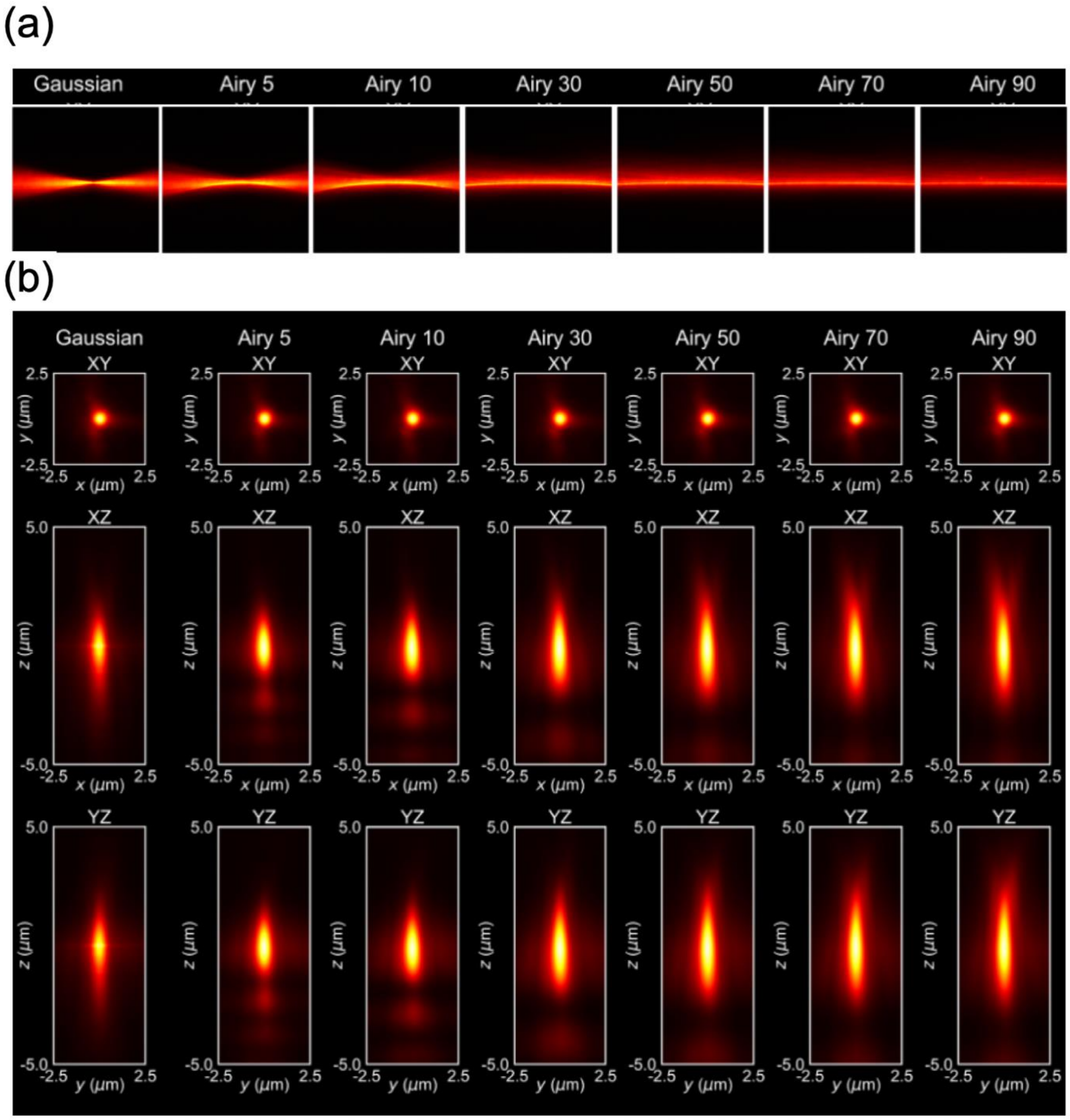


**Supplementary figure 2: Experimentally measured beam profiles and PSFs of Airy beams. (a)** Gaussian beam and Airy beams of strength 5, 10, 30, 50, 70, and 90, respectively. **(b)** PSF of Gaussian beam for comparison with Airy beams corresponding to phase masks of increasing strength (5, 10, 30, 50, 70, 90, respectively). *Top*: XY projections. *Middle*: XZ projections. *Bottom*: YZ projections. Obtained with 561 nm excitation wavelength and 100 nm diameter TetraSpeck beads suspended in 1.2% agarose in water.

### Supplementary figure 3:

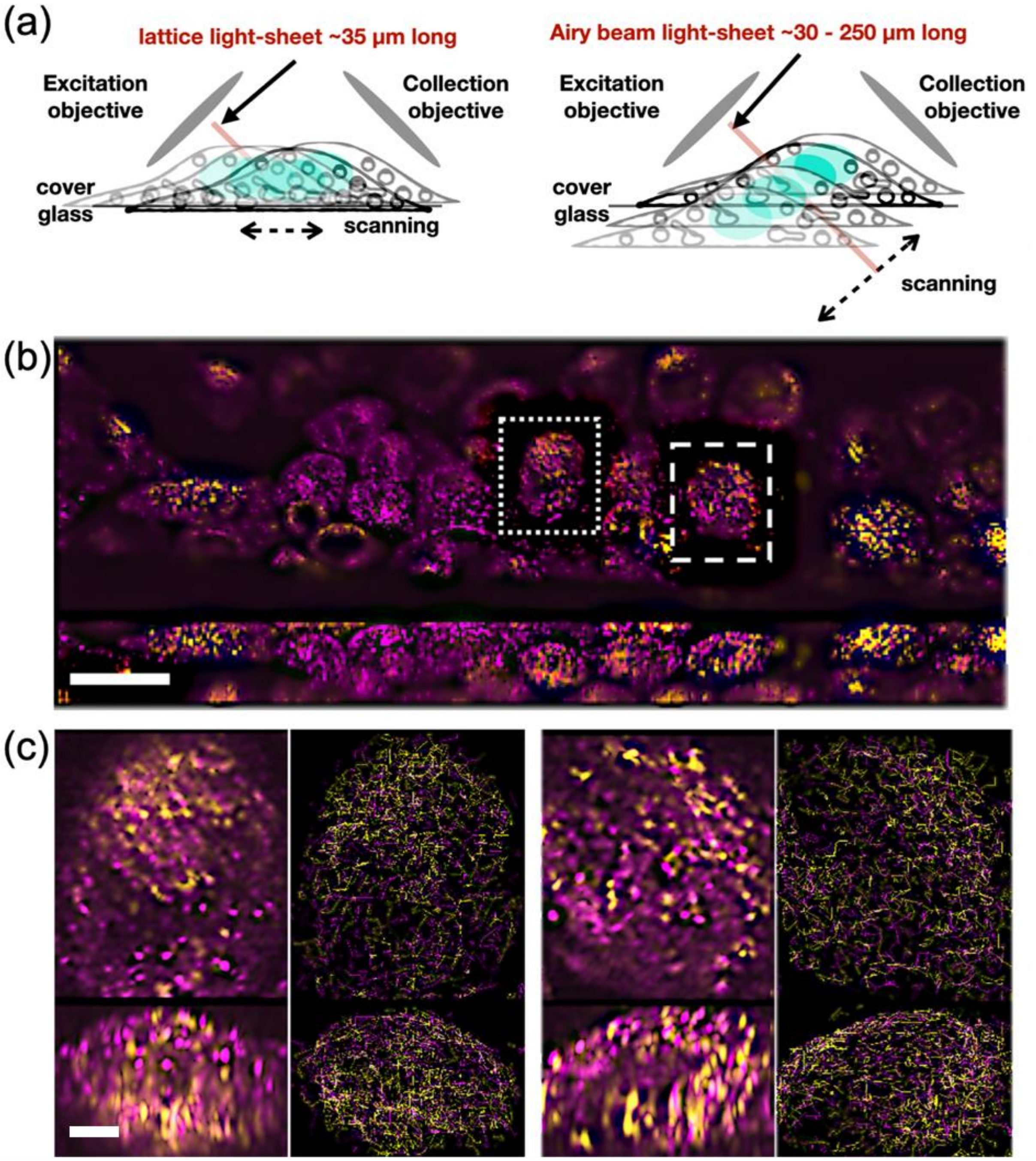


**Supplementary figure 3**: Simultaneous assessment of endosomal motility from many single cells. (a) Schematic comparing acquisition from lattice light-sheet (left) and Airy beam light-sheet (right) microscopes. (b) To analyse endosomal motility in single cells, dual-channel images were simultaneously acquired of HeLa cells expressing Rab5-GFP and Rab7-mCherry, spanning approximately 127 µm × 302 µm × 25 µm and imaged over 17 min at 5.7 s per volume. Max intensity projections taken from (top) XY plane and (bottom) YZ plane corresponding to (left) raw images and (right) tracks of Rab5-positive (magenta) and Rab7-positive (yellow) endosomes, showing endosomes tracked for at least 5 frames (30 s) during a representative time window of 5 min. (c–d) Selected regions corresponding to dotted (c) and dashed (d) insets in b. Panels are as described in **b**. Scale bars: **(b):** 25 µm, **(c):** 5 µm.

### Supplementary figure 4:

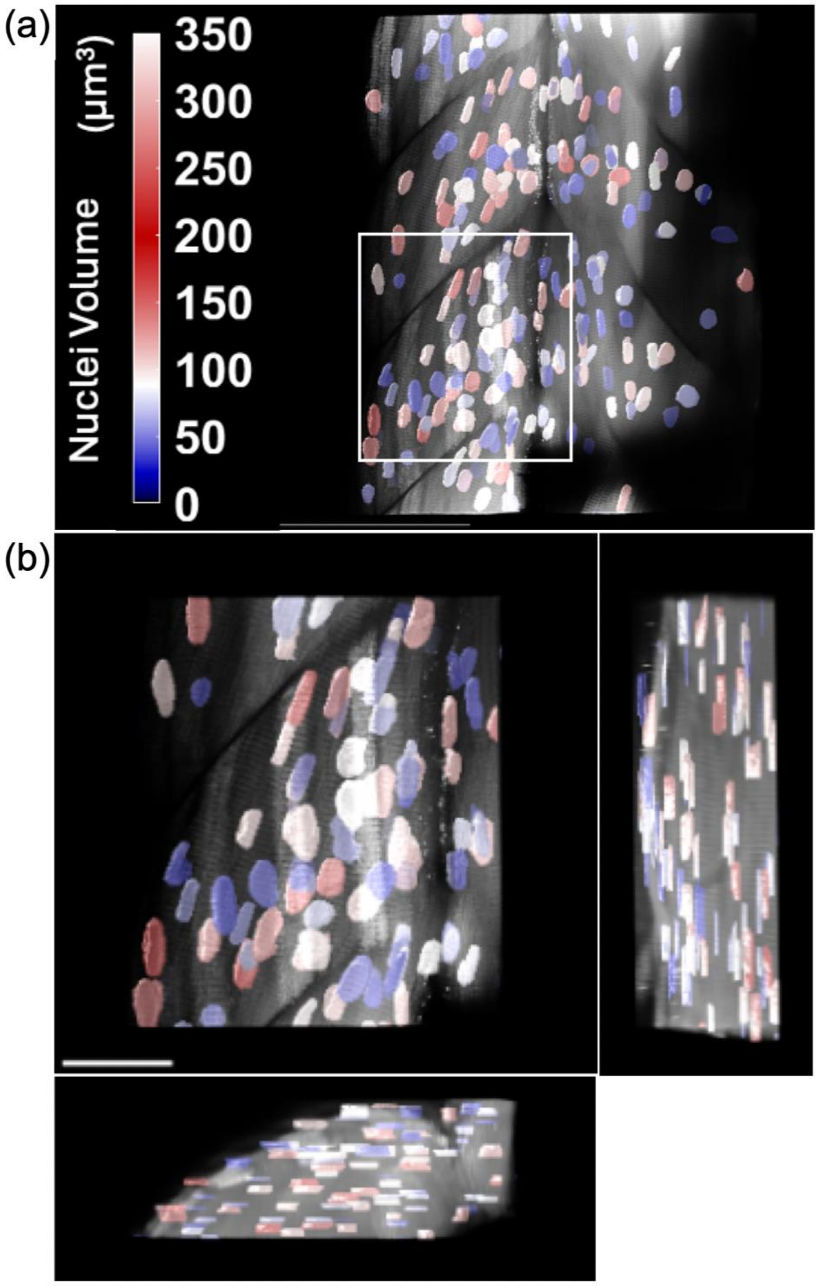


**Supplementary Figure 4: Morphometric analysis of nuclear architecture in zebrafish skeletal muscle. (a)** Representative maximum intensity projection of a 3 days post-fertilization (dpf) transgenic *Tg(actc1b:mito-GFP)*$^{uom407Tg}$; *Tg(actc1b:ER-mCherry)*$^{uom408Tg}$ larva. The endoplasmic reticulum (ER) channel is shown in greyscale, overlaid with nuclei segmented from the negative space in the ER channel, which are colour-coded by volume. **(b)** Zoomed orthogonal projections corresponding to the white bounding box indicated in **a**. Scale bars: **(a)** 100 µm, **(b)** 30 µm.

### Supplementary figure 5:

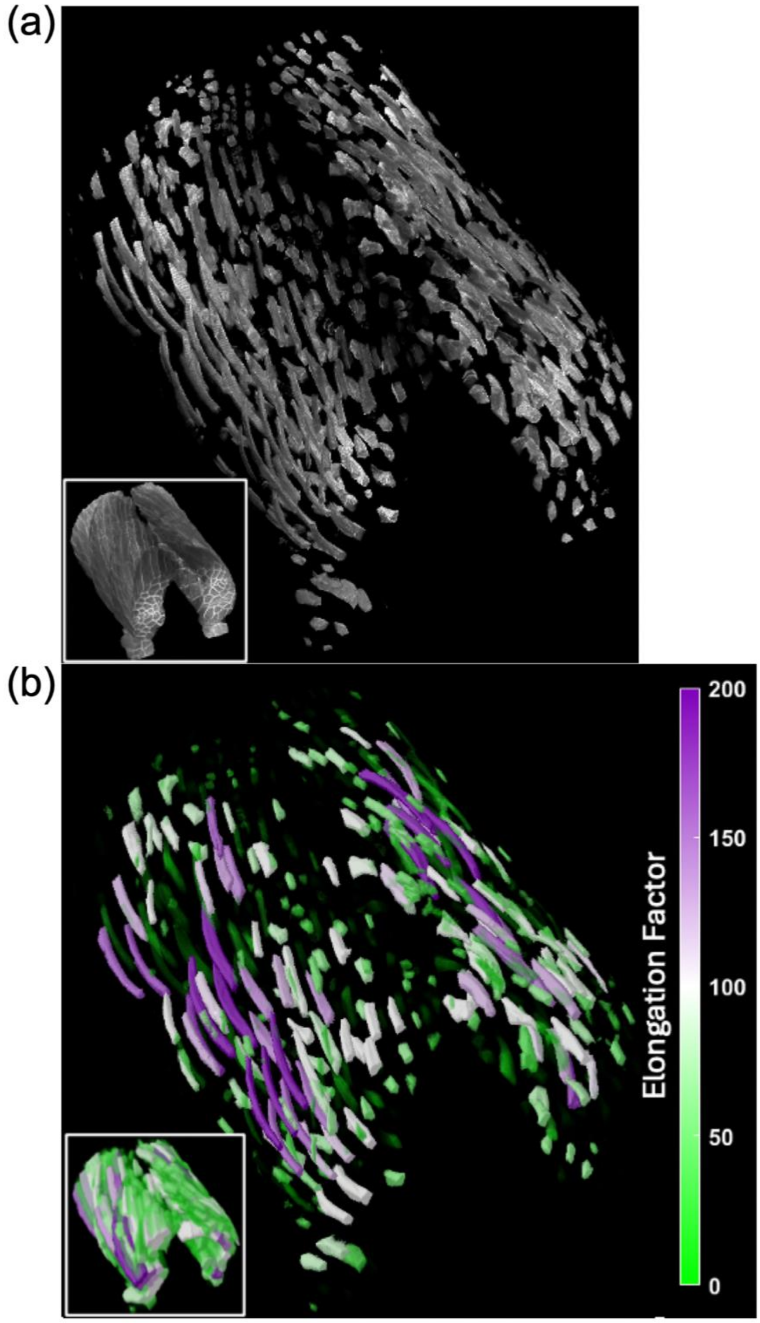


**Supplementary figure 5: Morphometric analysis of cellular architecture in zebrafish skeletal muscle. (a)** Raw volumetric data of a 3-day post-fertilization (dpf) transgenic *Tg(acta1:mCherryCAAX)*$^{pc22Tg}$ zebrafish larva, with muscle cells labelled with CaaX-mCherry, showing both computationally separated and unseparated (inset, bottom left) individual myofibers. **(b)** 3D segmented muscle cells colour-coded by their elongation factor, providing a quantitative measure of myofiber morphology. The accompanying colour bar maps the elongation factor values.

### Supplementary figure 6:

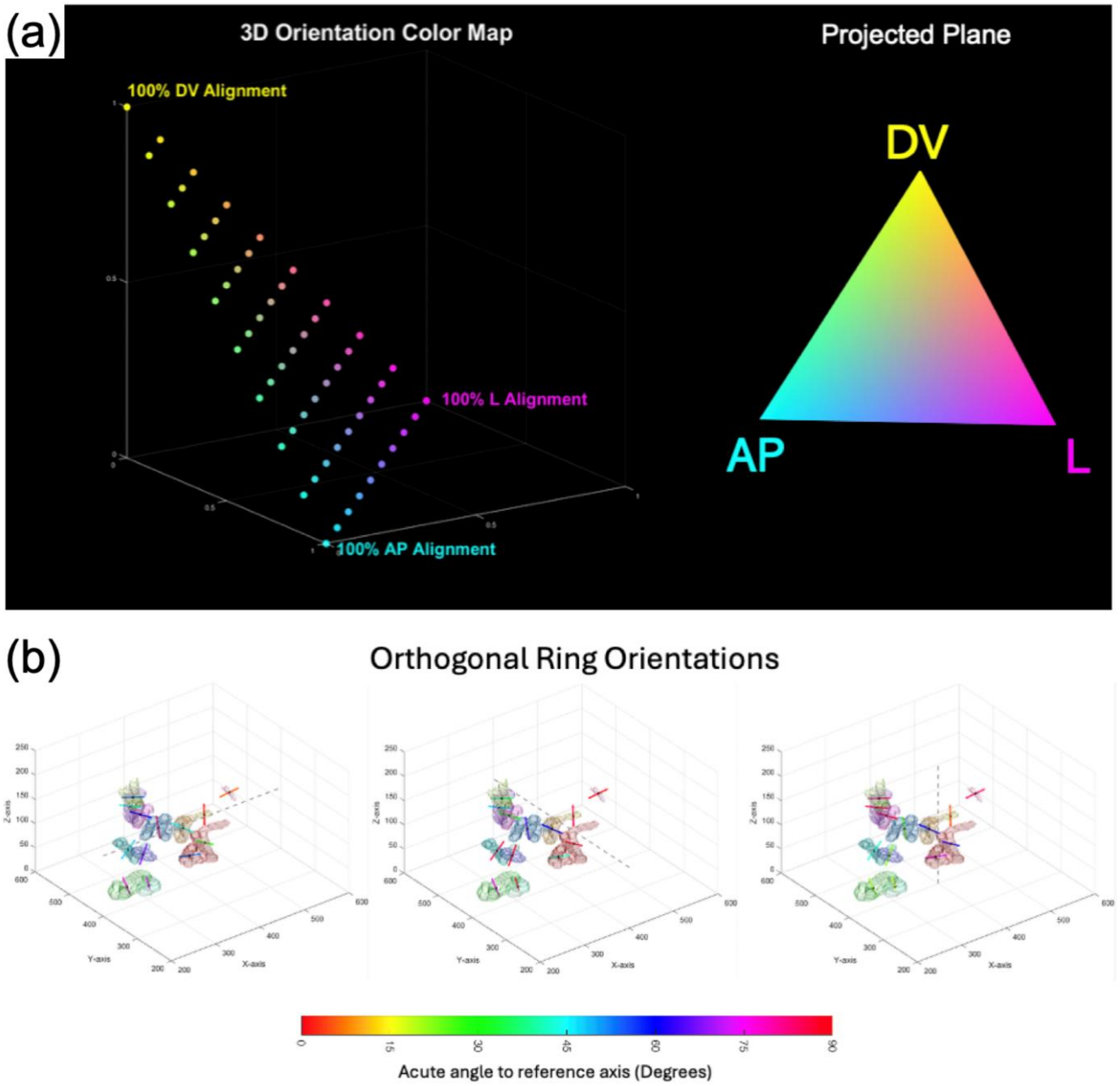


**Supplementary figure 6: 3D orientation mapping and quantitative analysis of annotated ring structures in Drosophila. (a)** Visualisation of the 3D Orientation Colour Map strategy. (*Left*) The vector endpoint colour map demonstrates the mapping of 3D spatial orientations based on the degree of alignment with the dorsal-ventral (DV), anterior-posterior (AP), and lateral (L) axes. (*Right*) A 2D projection of the colour plane situated at single positive unit distance from the origin, illustrating continuous color values assigned directional alignments. **(b)** Quantitative analysis of Orthogonal Ring Orientations. Surface renderings of a volumetric subsection display manually annotated ring structures. Three orthogonal reference axes (indicated by dashed grey lines) serve as the basis for orientation measurement. The rings are colour-coded based on the acute angle (0° to 90°) measured relative to each respective reference line, providing a spatial representation of structural alignment within the volume.

### Supplementary figure 7:

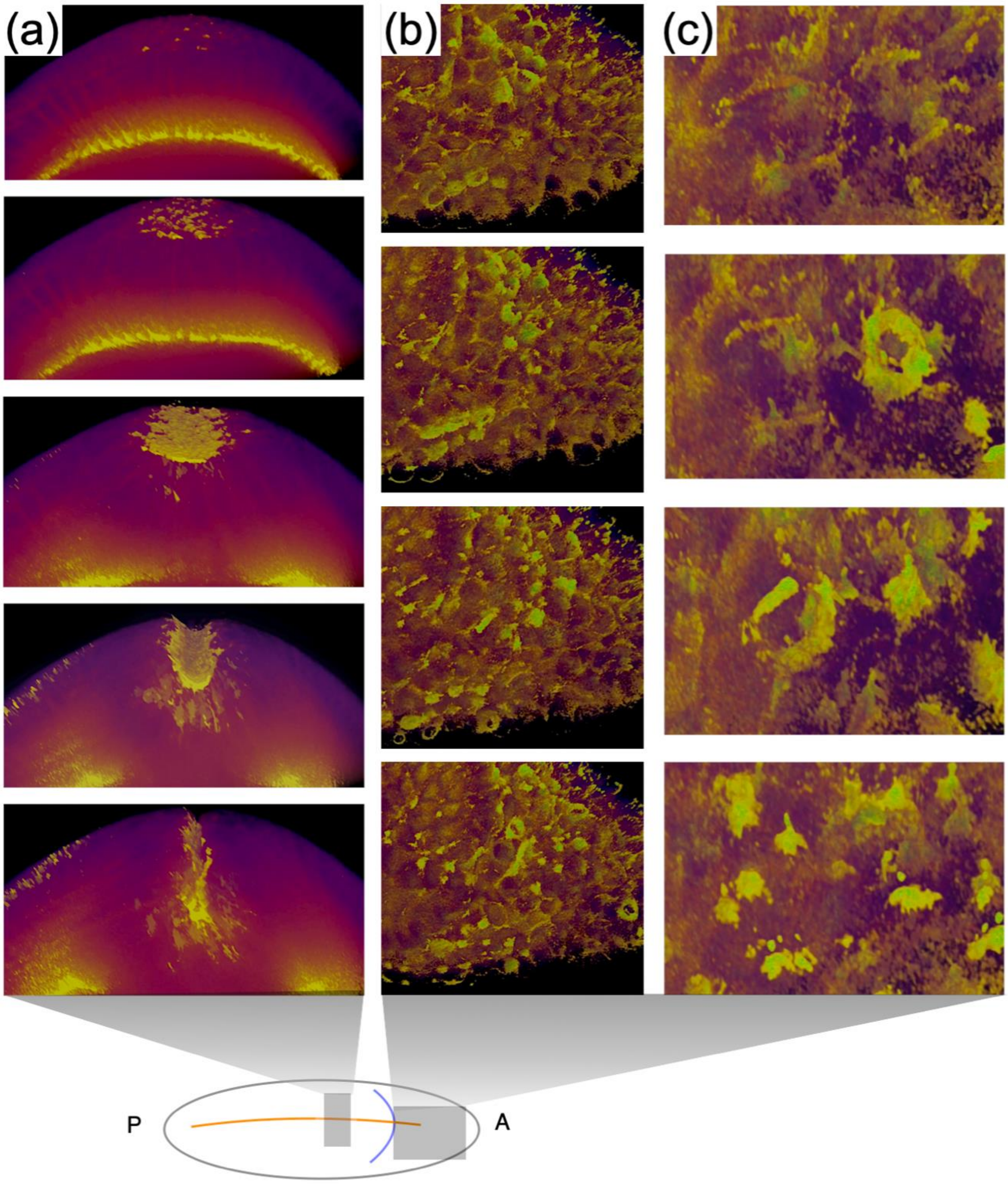


**Supplementary figure 7: Spatiotemporal dynamics of ventral furrow formation and cell divisions in the cephalic region. (a)** Cross-section view of a transgenic *Drosophila* embryo expressing s qh-3x-GFP during ventral furrow formation. The ventral side of the embryo is at the top of the FoV. **(b)** Cephalic region of the same embryo. **(c)** Detail-view of individual mitotic divisions within the cephalic region. **(d)** Schematic indicating depicted regions in **a** and **b** along the anterior-posterior axis of the embryo.

### Supplementary figure 8:

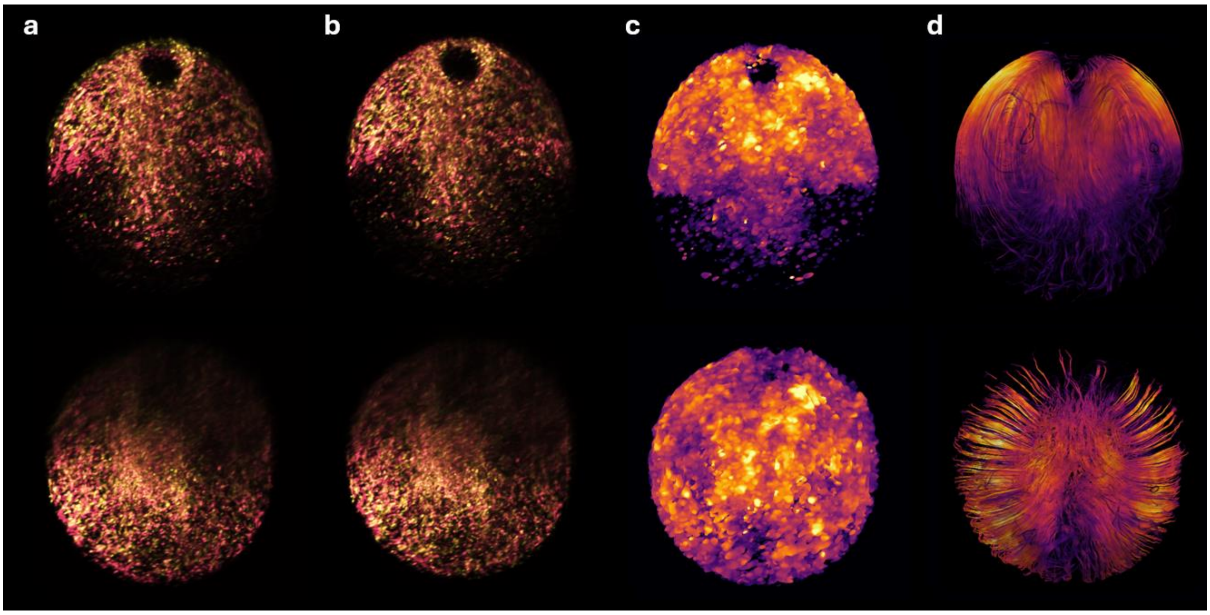


**Supplementary figure 8: Calculating 3D mitochondrial flow maps in mouse oocytes. (a)** Raw, unregistered images of the same oocyte taken from the current (*magenta*) and first (*yellow*) frames. Images separated by ~1 hour are shown to aid in visualisation of translational drift. The top view shows a hemisphere centred on the equator; the bottom shows a hemisphere centred on the north pole. **(b)** Images of the same oocyte as in **a**, after performing registration to correct for translational drift. **(c)** The optical flow between subsequent frames consists of three 3D stacks corresponding to the flow at each point, along each axis. The magnitude of flow at each point (i.e., excluding direction) is shown here as a single 3D stack. **(d)** To visualise flow velocity (i.e., including both magnitude and direction), 3D streamlines were calculated from the time-averaged flow map over an hour of imaging.

### Supplementary figure 9:

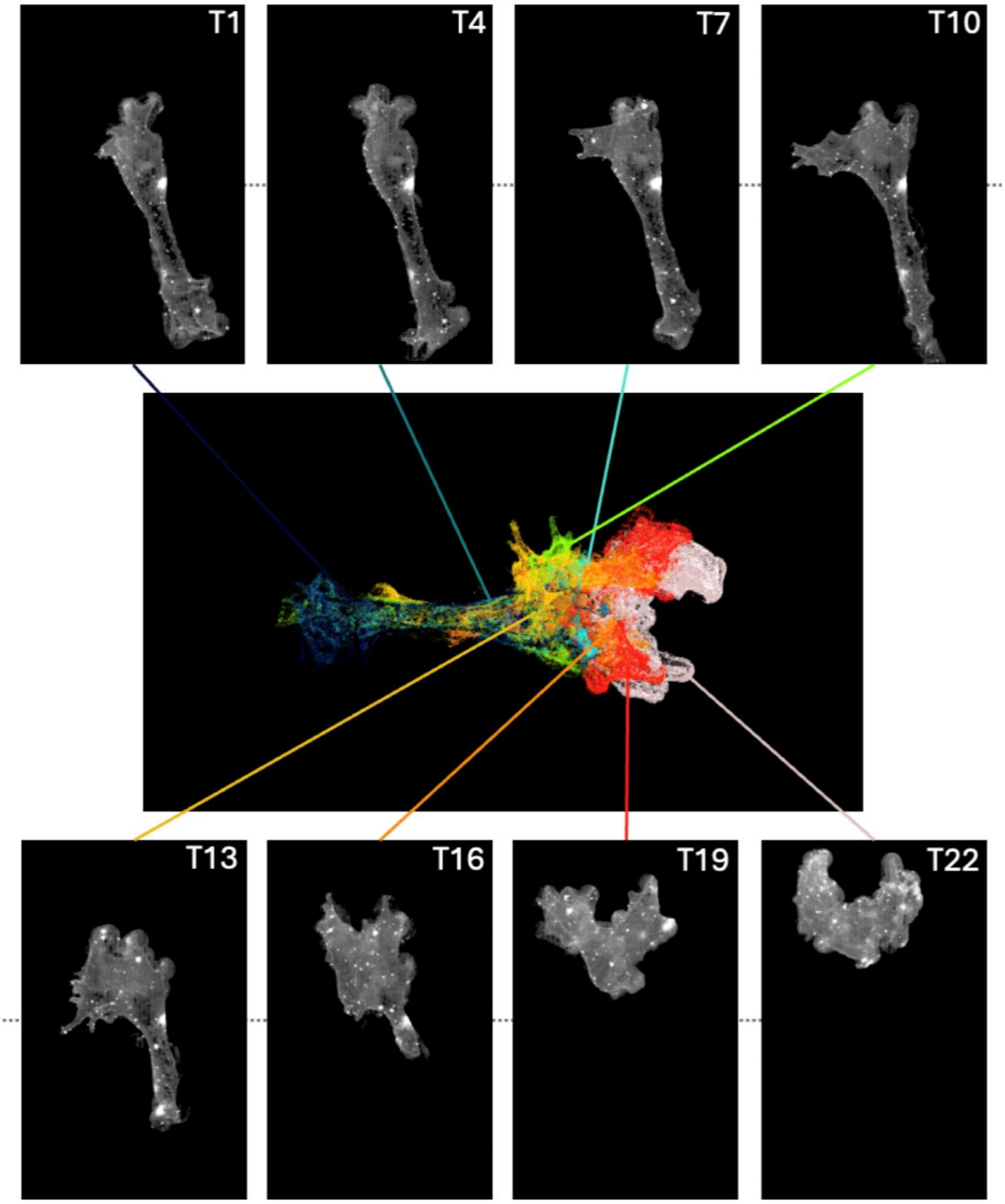


**Supplementary figure 9: Spatiotemporal dynamics of amoeboid morphological transitions.** Montage of representative frames from a high-speed volumetric time-series of an *Amoeba proteus* cell. Top and bottom panels display individual volumes at 18.6 s intervals (every third captured frame, T1 through T22). The central composite image illustrates the temporal colour-coding of the integrated dataset, where successive morphological changes are mapped to a specific colour spectrum. This projection highlights the progression of dynamic morphological changes. Data were acquired at a temporal resolution of 6.2 s per volume; intermediate frames between the displayed panels are omitted for visual clarity.

### Supplementary movie 1:

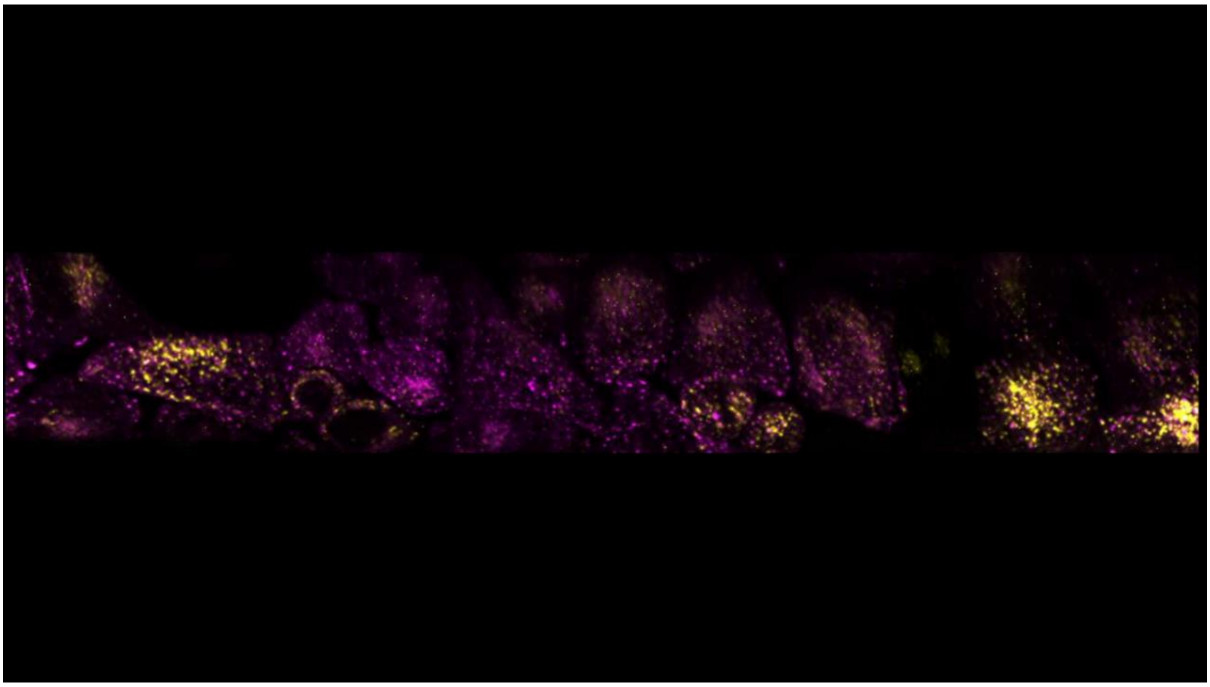

**Supplementary movie 1: Simultaneous assessment of endosomal motility from many single cells.** Dual-channel images were acquired simultaneously of HeLa cells expressing Rab5-GFP and Rab7-mCherry, spanning approximately 127 μm × 302 μm × 25 μm and imaged over 17 min at 5.7 s per volume. Trajectories of endosomes observed for at least 5 frames (30 s) are overlaid.

### Supplementary movie 2:

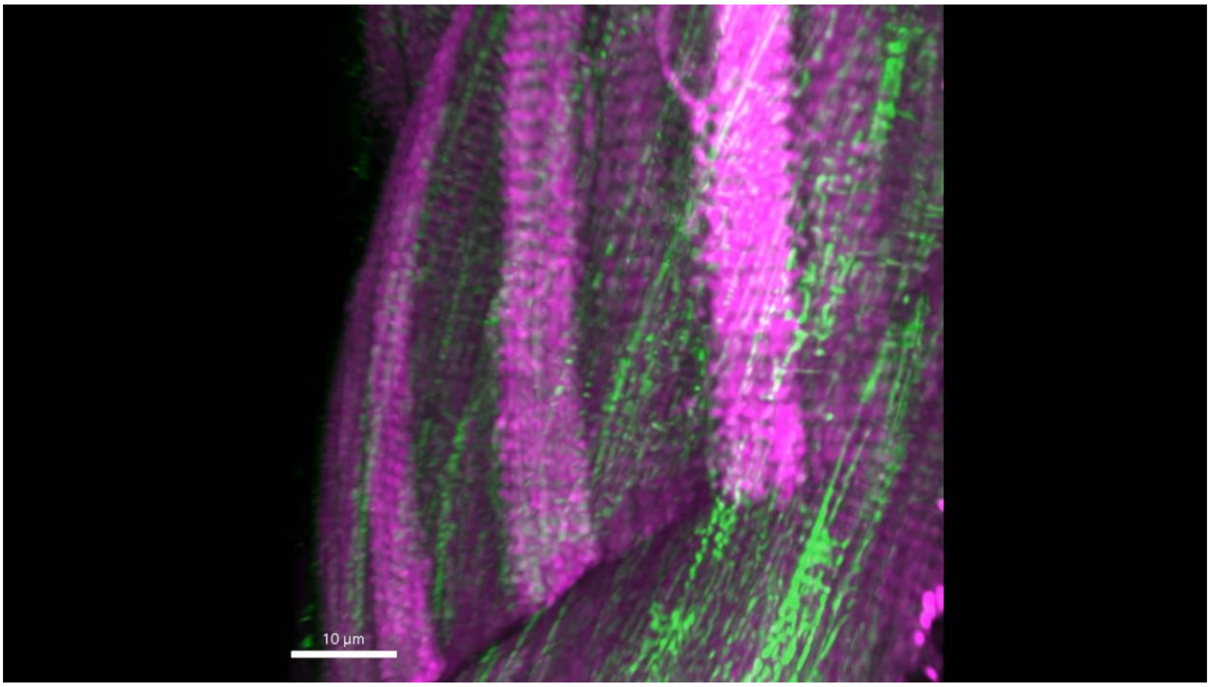


**Supplementary movie 2: Multiscale organisation in zebrafish skeletal muscle tissue.** Raw image of skeletal muscle tissue of a 3 dpf transgenic *Tg(actc1b:mito-GFP)*$^{uom407Tg}$ and *Tg(actc1b:ER-mCherry)*$^{uom408Tg}$ double positive larva at different zooms, showcasing different scales of organisation from the whole tissue to individual myotomes and mitochondria.

### Supplementary movie 3:

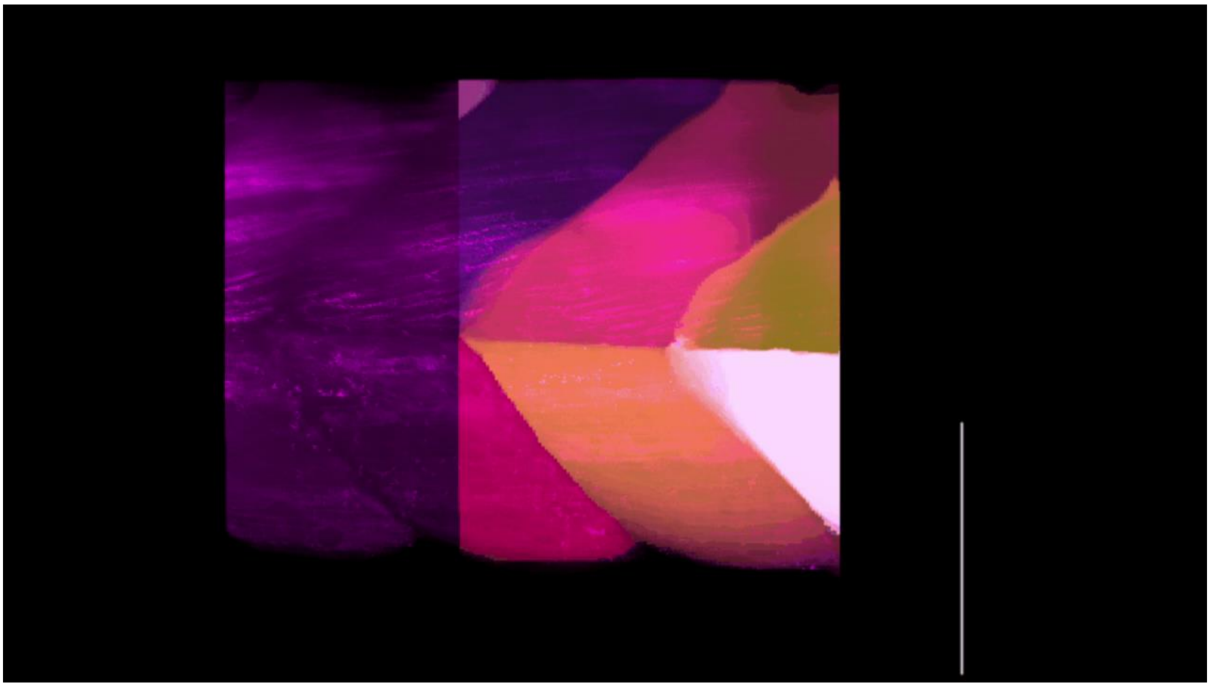

**Supplementary movie 3: Overview of analysis protocol of structures in zebrafish skeletal muscle tissue.** Raw image of skeletal muscle tissue of a 3 dpf transgenic *Tg(actc1b:mito-GFP)*[uom407Tg] and *Tg(actc1b:ER-mCherry)*[uom408Tg] double positive larva, illustrating 3D segmentations of individual myotomes, mitochondria, and nuclei.

### Supplementary movie 4:

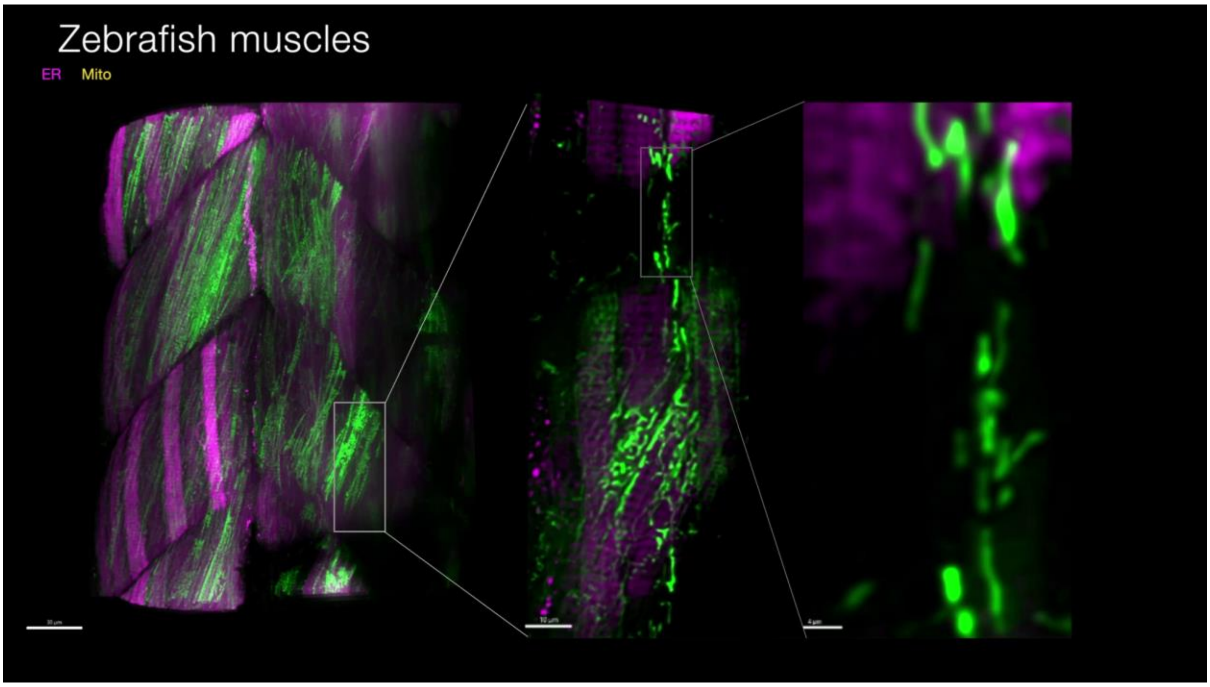


**Supplementary movie 4: Mitochondrial dynamics in large volumes of zebrafish tissue.** Movie of skeletal muscle tissue of a 3 dpf transgenic *Tg(actc1b:mito-GFP)*$^{uom407Tg}$ and *Tg(actc1b:ER-mCherry)*$^{uom408Tg}$ double positive larva at progressively higher zooms, showcasing mitochondrial dynamics over time.

### Supplementary movie 5:

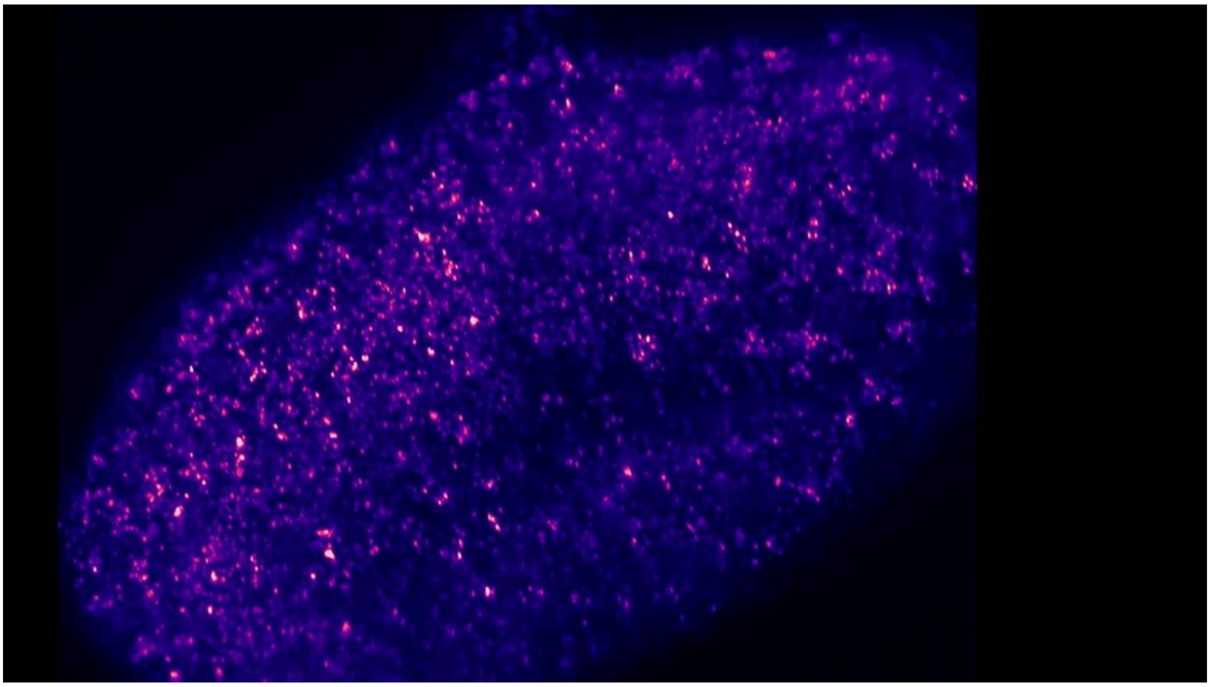

**Supplementary movie 5: Endosomal motility in zebrafish embryo tail tissue.** Raw movie of tailbud region of a 14-somite stage from a zebrafish embryo expressing mKate2-rab5ab after mRNA injection. The volume spans approximately 266 × 266 × 60 $\mu m^3$ and was imaged over 12 min at 5.9 s per volume.

### Supplementary movie 6:

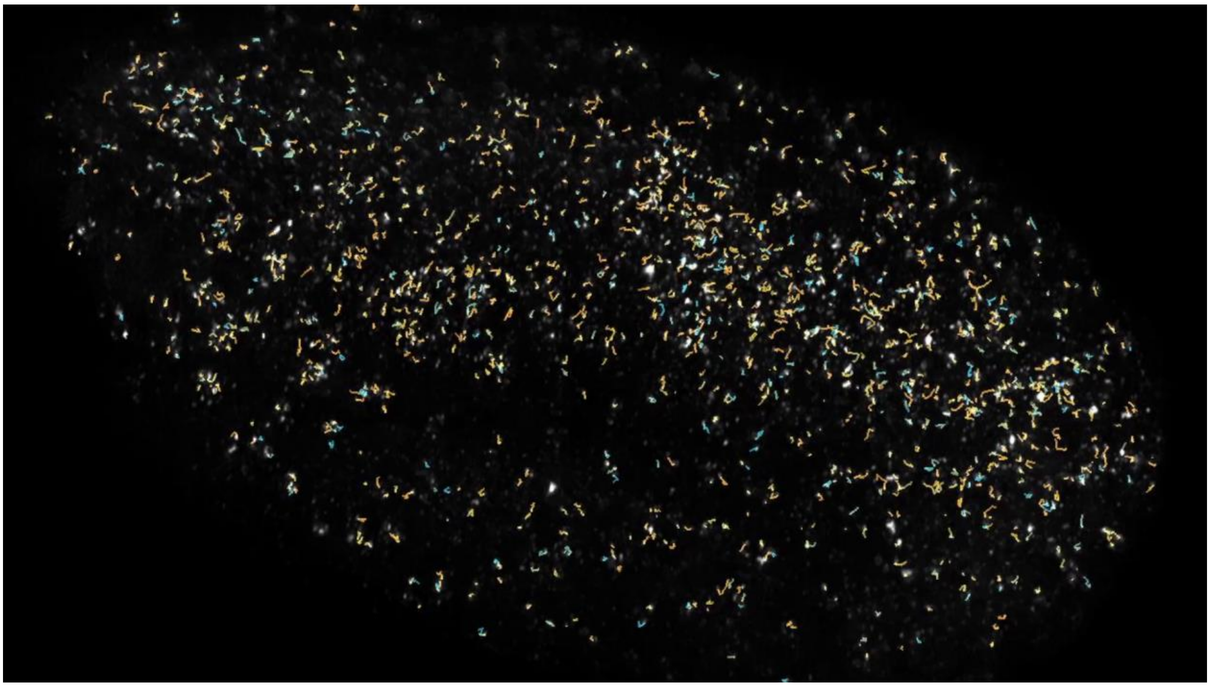

**Supplementary movie 6: Endosomal tracks in zebrafish embryo tail tissue.** Raw movie of the tailbud region of a 14-somite stage from a zebrafish embryo expressing mKate2-rab5ab after mRNA injection, overlaid with trajectories of mKate2-rab5ab-labelled endosomes tracked for at least 10 frames (60 s). Track segments are coloured according to the anomalous diffusion exponent (α) calculated by fitting the mean-squared displacement (MSD) for each segment. The volume spans approximately 266 × 266 × 60 $\mu m^3$ and was imaged over 12 min at 5.9 s per volume.

### Supplementary movie 7:

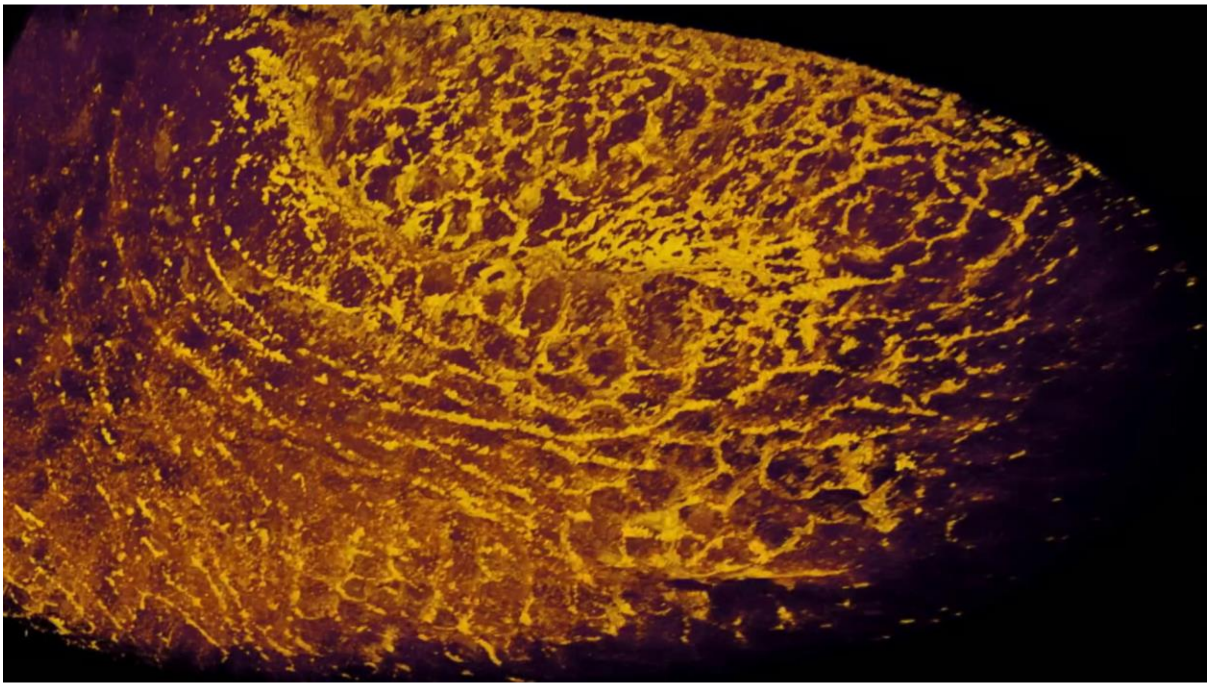

**Supplementary movie 7: Posterior midgut invagination in *Drosophila*.** Side view of posterior midgut (PMG) invagination at the posterior end of a *Drosophila* embryo expressing sqh-3x-GFP. Cells appear to turn and elongate with the progress of invagination, subsequently undergoing cell divisions, inferred by the formation of cytokinetic rings.

### Supplementary movie 8:

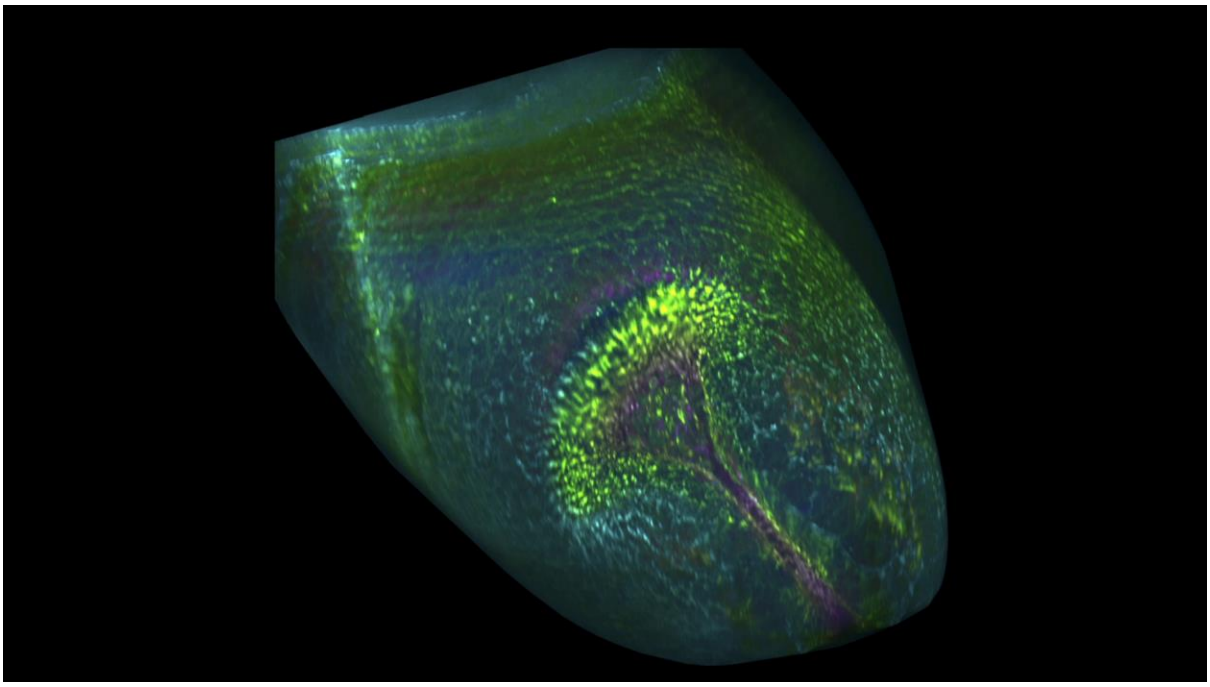

**Supplementary movie 8: Germ band elongation in Drosophila.** Movie of germ band elongation (GBE) in a *Drosophila* embryo expressing sqh-3x-GFP. Images are colour-coded according to distance from the embryo surface, ranging from yellow (superficial) to cyan (intermediate) to magenta (deep).

### Supplementary movie 9:

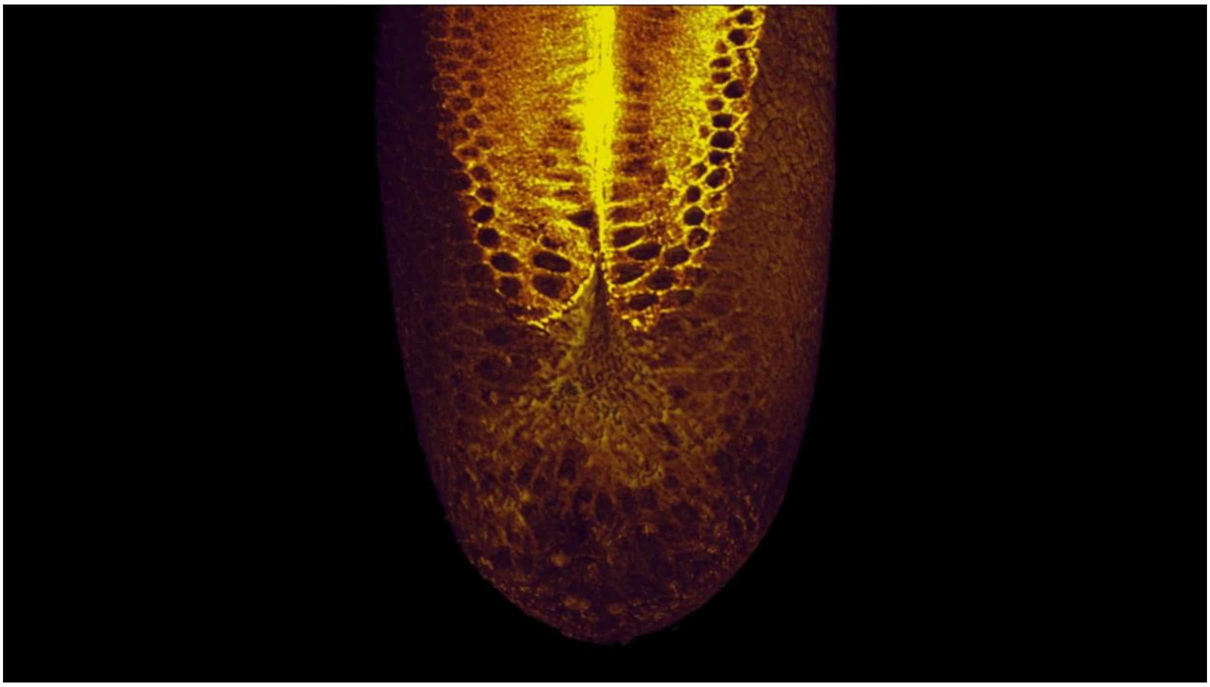

**Supplementary movie 9: Ventral slice through *Drosophila* embryo following posterior midgut invagination.** Top view of *Drosophila* embryo expressing sqh-3x-GFP, showing a cross-section at a depth of 35 μm. Cell divisions are visible as cytokinetic rings interspersed throughout the movie.

### Supplementary movie 10:

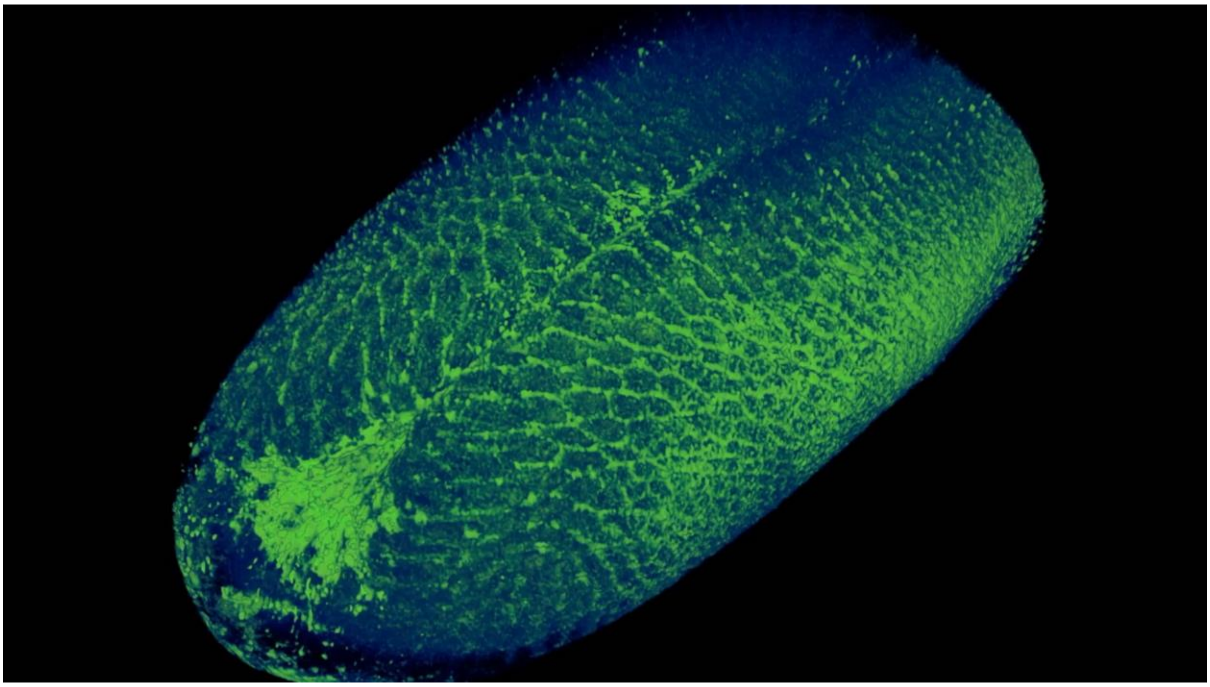

**Supplementary movie 10: Germ band extension in the posterior-ventral side followed by mitotic waves.** Patterns of divisions, along the anterior-posterior length, occur on the posterior-ventral side of the embryo, as the rapid phase of GBE concludes.

### Supplementary movie 11:

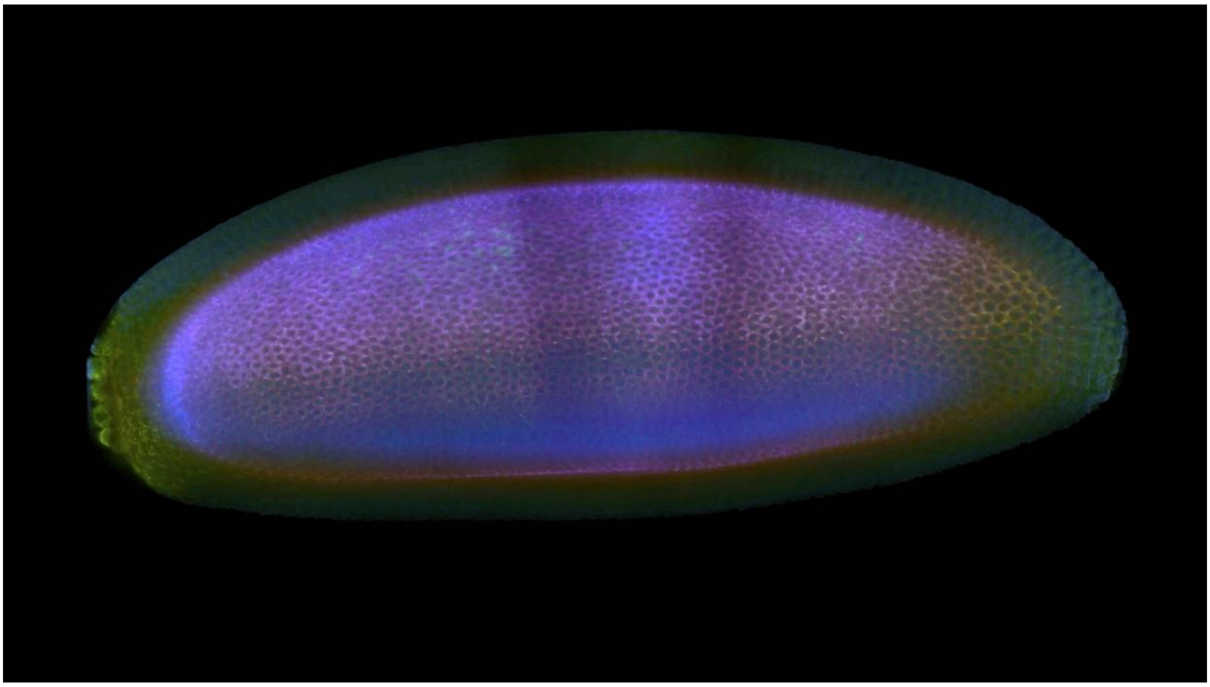

**Supplementary movie 11: Whole *Drosophila* embryo.** Two tiles of a *Drosophila* embryo expressing sqh-3x-GFP were stitched together to generate a movie of the entire embryo, taken at 1 frame every 66 s . Images are colour-coded according to distance from the embryo surface, ranging from yellow (superficial) to cyan (intermediate) to magenta (deep).

### Supplementary movie 12:

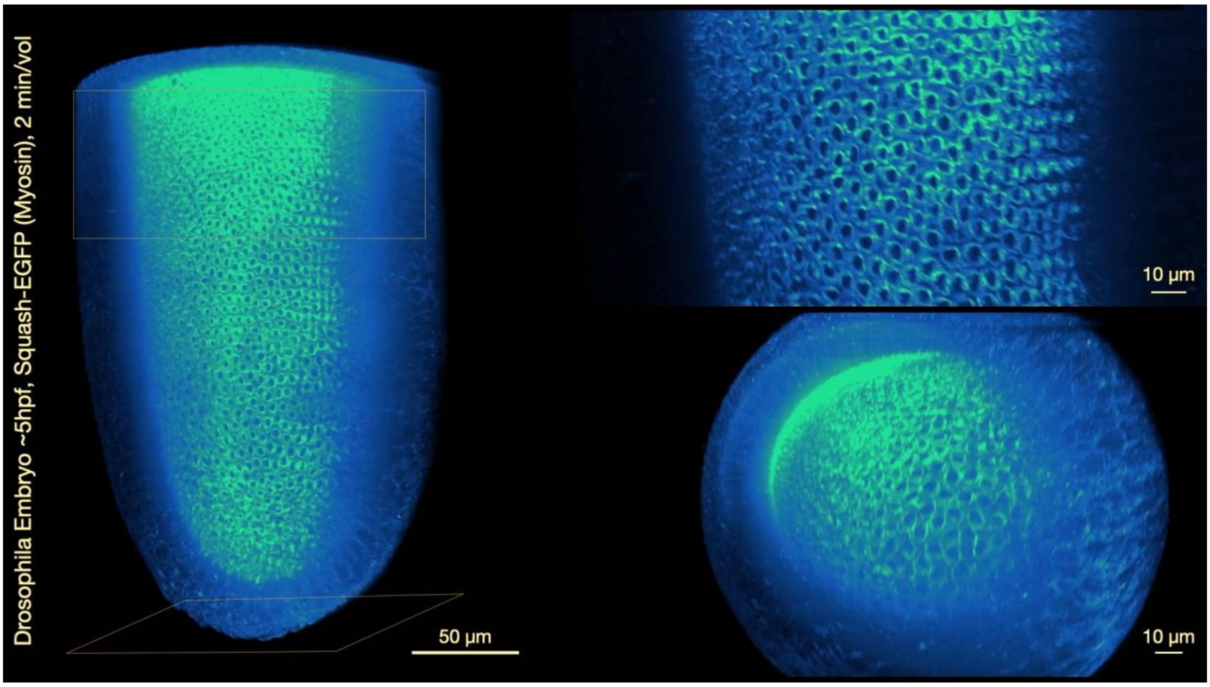


**Supplementary movie 12: sqh-GFP in *Drosophila* embryo reveals first mitotic waves post cellularis ation**. Waves of division in the precephalic region, which occur concurrently with the formation of cell boundaries after ventral furrow formation, can also be mapped.

### Supplementary movie 13:

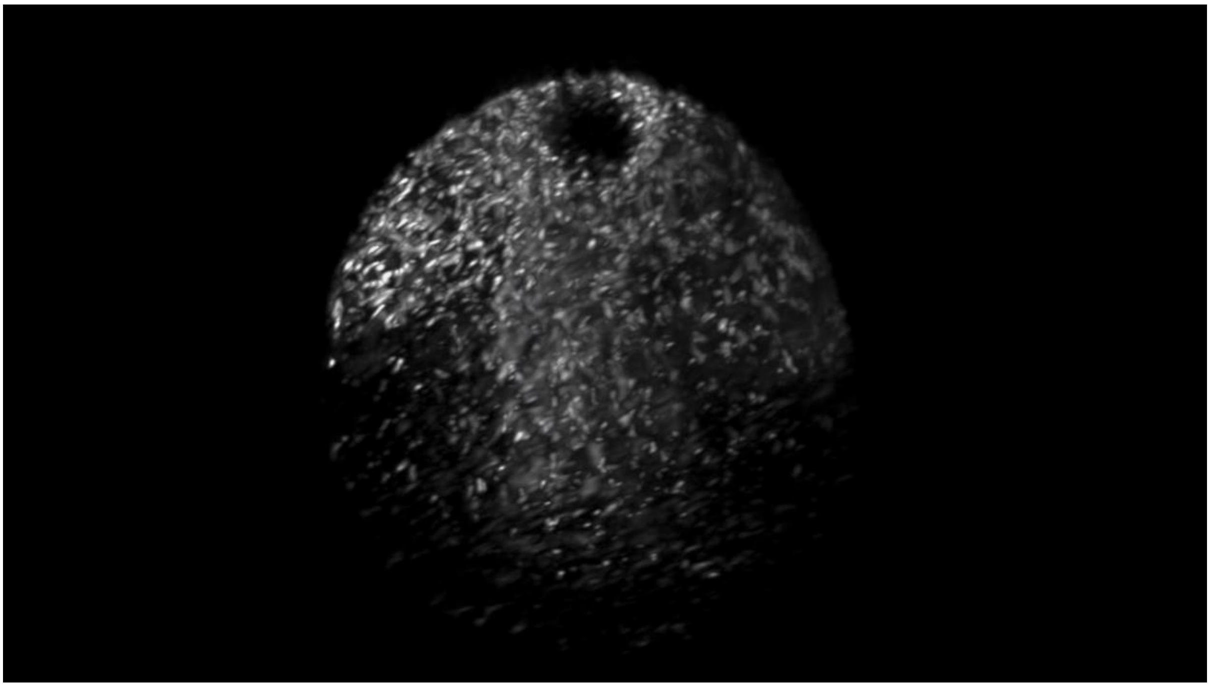

**Supplementary movie 13: Mitochondrial flow in a mouse oocyte.** Mouse meiosis II oocyte with labelled mitochondria (mito-Dendra2), with a diameter of ~80 µm was imaged every 5 min for 4 h, displaying large-scale mitochondrial flow throughout the cytoplasm.

### Supplementary movie 14:

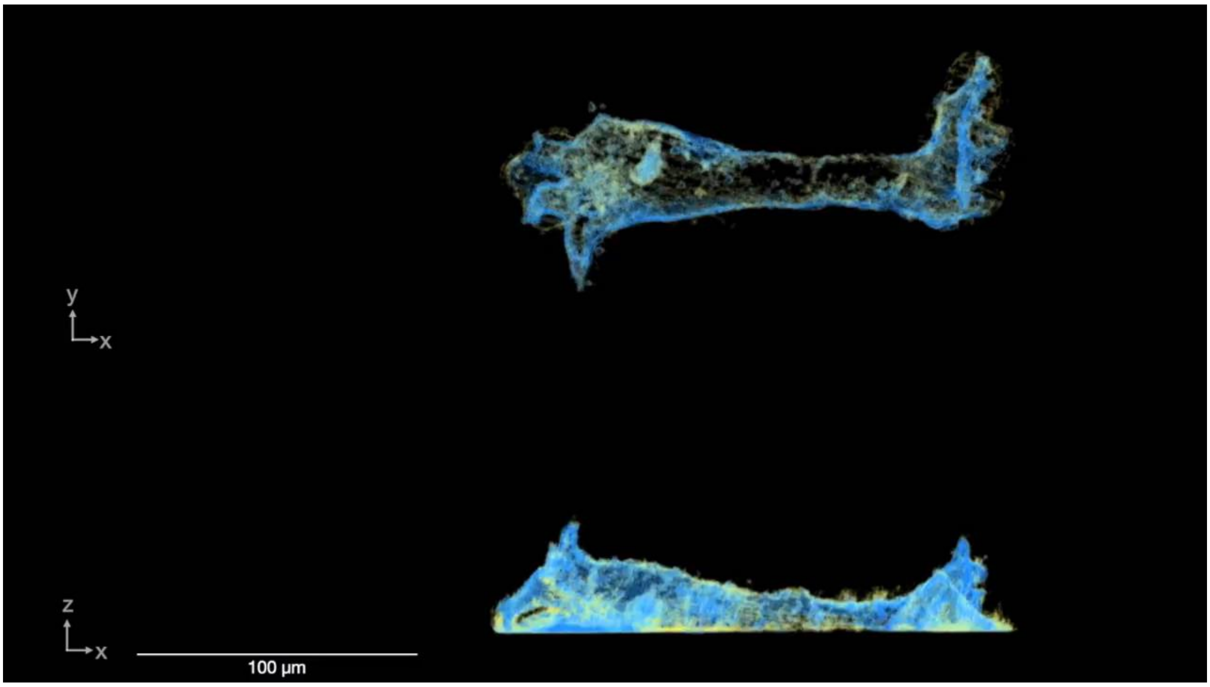


**Supplementary movie 14: Large-scale, rapid movement of an *Amoeba proteus* cell.** Movie of an amoeba captured at 6.2 s per volume.

### Supplementary movie 15:

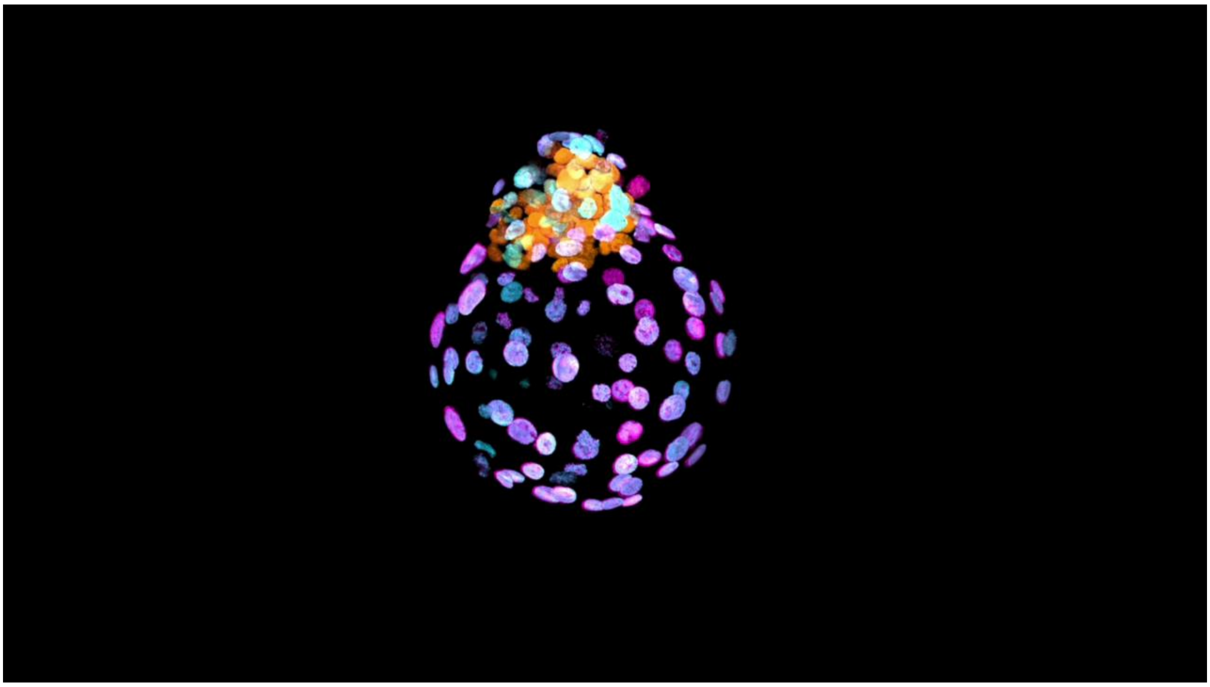

**Supplementary movie 15: Flythrough of a segmented iBlastoid.** Raw images of a representative iBlastoid stained for GATA3 (magenta), Nanog (orange), and GATA6 (cyan) visualised from different angles and slices, as well as instance segmentation of individual nuclei.
