## supplementary tables for "Observing concurrent subcellular dynamics in large living tissues"

[†] Equal first authors

#### Supplementary table T1

| Technique | Type of Beam | Datasets | Excitation Objective Used | Emission Objective Used | xyz PSF | xyz Claimed Resolution | Sample Demonstration | Temporal Resolution | Single-Plane FOV/Captured Volume | FOI/Depth/Propagation Features |
|---|---|---|---|---|---|---|---|---|---|---|
| SPIM(*1*) | Gaussian | Data1 (Highest Spatial Resolution) - Fig. S4 | Optical fiber + Cylindrical lens | 40x/0.8 | Not shown | Lateral: "well below 1µm" | Individual cell membranes and pole cells of Drosophila embryo | NA | 217µm×165µm | Light sheet thickness ranges from 3–10 µm. Using a 10×/0.30 objective, the beam waist reaches 6 µm with <42% width variation across a 660 µm FOV. |
| " | " | Data2 (Fastest Temporal Resolution) - Movie S4 | Optical fiber + Cylindrical lens | 5x/0.25 | Not shown | Lateral: 1.1um Axial: "About 6um" | Beating heart of a Medaka embryo | 10.7 planes/sec | 1500um×850um | " |
| " | " | Data3 (Largest FOV) - Fig. 2, S2 | Optical fiber + Cylindrical lens | 5x/0.25 | Not shown | Lateral: 1.1um | Medaka embryo | 1-4 planes/sec | 1500um×850um | Multiview reconstruction of four stacks from different |

| | | | | | | | | | | |
|---|---|---|---|---|---|---|---|---|---|---|
| | | | | | | Axial: "About 6um" | | | | orientations produced a complete 1.5 mm × 0.9 mm sample volume. |
| DLSM-SI(*2*) | Scanned Gaussian | Data1 (Highest Spatial Resolution) - Fig. 3 | 5x/0.16 | 20x/1.0 | Not shown | No Claim | Drosophila embryo nuclei | 2 SI planes/sec X4 (Multiview) | 757µm×757µm | DSLM-SI imaging of stage 6 *Drosophila* embryos achieves an average 50% signal decay penetration depth of 38.0 ± 3.8 µm. |
| " | " | Data2 (Fastest Temporal Resolution) - Fig. 2 | 5x/0.16 | 10x/0.45 | Not shown | No Claim | Medaka embryo membrane and nuclei | 6 DSLM planes/sec<br>2 SI planes/sec | 1516µm×1516µm | Captured planes represent the entire embryo to a detection depth of ~300 µm, covering roughly 25% of the central yolk cell surface. DSLM-SI achieves full penetration of one-day-old embryo head structures, including the forebrain, midbrain, and eyes. |
| " | " | Data3 (Largest FOV) - Fig. 2 | 5x/0.16 | 10x/0.45 | Not shown | No Claim | Medaka embryo membrane and nuclei | 6 DSLM planes/sec<br>2 SI planes/sec | 1516µm×1516µm | " |
| Bessel Light Sheet(*3*) | Bessel | Data1 (Highest Spatial Resolution) Fig. S8 (Bessel SI 9 Phases) | 40x/0.8 | 40x/0.8 | Axial: 0.29um<br>Lateral: ~0.3um (Abbe limit) | ~0.3um 3D isotropic | 100nm Beads | NA | NA | Scanned Bessel beams, combined with structured illumination or two-photon excitation, generate light sheets thinner than 0.5 µm for high-resolution 3D subcellular imaging. |
| " | " | Data2 (Fastest Temporal Resolution) Fig. 5 (Bessel TPE) | 40x/0.8 | 40x/0.8 | Axial: 0.49um | No claim | Histone H2B in an LLC-PK1 cell undergoing mitosis | 200 planes/sec | 33umx33um | Two-photon excitation (TPE) suppresses Bessel side lobes, enabling a 0.5 µm FWHM light sheet with a continuously swept beam. |
| " | " | Data3 (Largest FOV) Fig. 2e (Bessel TPE) | 40x/0.8 | 40x/0.8 | Axial: 0.49um | No claim | Fixed mitochondria | 12.5 planes/sec | 53umx80um | " |
| MuVi SPIM(*4*) | Gaussian | Data1 (Highest Spatial Resolution) Fig. 2e | 10x/0.3 | 40x/0.8 | Not shown | Lateral: 0.43um<br>Axial: 2.28um | Drosophila embryo membrane marker Gap43-mCherry | 43 seconds for entire embryo with 4 view reconstruction and 1 rotation 41ms exposure x2 (each arm) x2 (each view)<br>6 planes/sec | NA | Light-sheet thickness maintains a range of 2.3–3.2 µm across the entire 200 µm diameter of the *Drosophila* embryo. |
| " | " | Data2 (Fastest Temporal Resolution) Fig. 2 | 10x/0.3 | 25x/1.1 | Not shown | No claim | Drosophila embryo nuclei marker H2Av-mCherry | 10 seconds for entire embryo | 519umx438um (Inferred). At this magnification, the image of a typical Drosophila embryo optimally covers the sCMOS image sensor. | " |
| " | " | Data3 (Largest FOV) Fig. 2 | 10x/0.3 | 25x/1.1 | Not shown | No claim | Drosophila embryo nuclei marker H2Av-mCherry | 10 seconds for entire embryo | 519umx438um (Inferred). At this magnification, the image of a typical Drosophila embryo optimally covers the sCMOS image sensor. | " |
| SiMView(*5*) | Gaussian | Data1 (Highest Spatial Resolution) Fig. S4 | 4x/0.28 | Single photon: 40x/1.0 | Lateral (single photon): ~0.4um<br>Axial (single photon): 1.59um | 'Subcellular resolution' | 50nm beads<br>Nuclei, filopodial dynamics | 15s per timepoint at 30spv | ~410umx346um (Inferred) | Four complementary optical views provide near-complete physical coverage with a maximum 20 ms time shift, regardless of specimen size. |

| | | | | | | | | | | |
|---|---|---|---|---|---|---|---|---|---|---|
| " | " | Data2 (Fastest Temporal Resolution) Fig. 5 | 4x/0.28 | 16x/0.8 | Lateral (two photon): ~0.6um Axial (two photon): 1.87um | No claim | Drosophila syncytial blastoderm nuclei | 10s per timepoint at 25spv | ~1050umx886um (Inferred) | " |
| " | " | Data3 (Largest FOV) Fig. 3 | 4x/0.28 | 16x/0.8 | Lateral (two photon): ~0.6um Axial (two photon): 1.87um | No claim | Stage 16 drosophila embryo | 15s per timepoint at 30spv | ~1050umx886um (Inferred) | " |
| LLSM(*6*) | Lattice/Bessel | Data1(Highest Spatial Resolution) Fig 2. E, G | 20x/0.65 | 25x/1.1 | Lateral (SIM): 150nm Axial (SIM):280nm | Subcellular | According to OTF Beads taken but FWHM not quantified | NA | NA | Performance declines with depth due to sample-induced aberrations; imaging beyond 20–100 µm typically would require adaptive optics, depending on the specimen's optical heterogeneity. |
| " | " | Data2 (Fastest Temporal Resolution) Fig. 4C | 20x/0.65 | 25x/1.1 | Lateral: 230nm Axial: 370nm | Subcellular | Tetrahymena GFP-Scramblase | 3.2 volumes per second | 53umx53umx151um | |
| " | " | Data3 (Largest FOV) Fig. 6C | 20x/0.65 | 25x/1.1 | Lateral: 230nm Axial: 370nm | Subcellular | Drosophila eGFP-sGMCA mCherry-sqs | 8 per volume | 106umx53umx52um | |
| Airy beam(*7*) | Airy | Data1(Highest Spatial Resolution) Fig. 2c | 20x/0.42 | 20x/0.40 | Lateral: 1.9um Axial: <1.5um | Axial: 865nm | Fluorescent microspheres in PDMS | NA | 237umx178um (Inferred) | Axial FWHM remained < 1.5 µm over a 100 µm range |
| " | " | Data2(Fastest Temporal Resolution) Supp Fig. 2 | 40x/0.80 | 40x/0.80 | NA | NA | ACHN cell cluster / spheroid | NA | 118umx89um (Inferred) | The Airy light sheet can obtain high resolution throughout the 80-µm FOV |
| " | " | Data3(Largest FOV) Fig. 3 | 20x/0.42 | 20x/0.40 | Lateral: 1.9um Axial: <1.5um | Axial: 865nm | Nuclei in tail of juvenile amphioxus | NA | 237umx178um (Inferred) | |
| IsoView(*8*) | Gaussian | Data1(Highest Spatial Resolution) Sup Fig. 6c | Custom f15mm/0.714 | Custom f15mm/0.714 | Lateral: 0.6um Axial: 2.98um Multiview decon: (0.42um isotropic) | NA | Beads | NA | NA | In superficial tissues, IsoView improves PSF dimensions: lateral-vertical (1.77 to 1.09 µm), lateral-horizontal (2.06 to 1.68 µm), and axial (5.45 to 1.59 µm). Centrally, PSF improvements reach 1.20 µm (LV), 1.73 µm (LH), and 2.13 µm (axial). Average resolution remains 1.1–1.7 µm across most of the embryo, with deep, optically inaccessible regions maintained ≤2.5 µm. |
| " | " | Data2(Fastest Temporal Resolution) Fig. 2 | Custom f15mm/0.714 | Custom f15mm/0.714 | Lateral: 0.6um Axial: 2.98um Multiview decon: (0.42um isotropic) | 1.1-2.5um isotropic | Stage 17 drosophila embryos GCaMP6s | 2 volumes per second | NA | " |
| " | " | Data3(Largest FOV) Fig. 2 | Custom f15mm/0.714 | Custom f15mm/0.714 | Lateral: 0.6um Axial: 2.98um Multiview decon: (0.42um isotropic) | 1.1-2.5um isotropic | Stage 17 drosophila embryos GCaMP6s | 2 volumes per second | NA | " |
| AO-LLSM(*9*) | Lattice | Data1(Highest Spatial Resolution) Fig. 1e | 20x/0.65NA | 25x/1.1NA | NA | Diffraction-limited performance | tagRFPt-Clathrin LCA EGFP-Dynamin in Organoid hESCs | 2.8s per volume | 49.7umx68.3umx30.2um | 30 µm beam length utilized. |
| " | " | Data2(Fastest Temporal Resolution) Fig. 1e | 20x/0.65NA | 25x/1.1NA | NA | Diffraction-limited performance | tagRFPt-Clathrin LCA EGFP-Dynamin in Organoid hESCs | 2.8s per volume | 49.7umx68.3umx30.2um | 30 µm beam length utilized. |

| | | | | | | | | | | |
|---|---|---|---|---|---|---|---|---|---|---|
| " | " | Data3(Largest FOV) Fig. 4a,b,c | 20x/0.65NA | 25x/1.1NA | NA | Diffraction-limited performance | EGFP-membrane Zebrafish 96hpf | 11.1min per volume | 213umx213umx113um (stitched) | 30 μm beam length utilized with a 7×7×3 volume stitching configuration. |
| Reflective Light-sheet(10) | Gaussian | Data1(Highest Spatial Resolution) Fig 4.b,c | 28.6x/0.71NA | 25x/1.1NA | NA | Lateral: 0.26um Axial: 0.28um (Four views) | According to OTF. | NA | Can be up to 110umx80um | NA |
| " | " | Data2(Fastest Temporal Resolution) Fig 2a, Movie 3 | 40x/0.8NA | 40x/0.8NA | NA | 0.33um isotropic | Nematode embryo expressing GCaMP3 | 2.86 volumes per second | 40umx60um | No significant deterioration in image quality was observed in modestly sized (thickness <50 μm) samples. |
| " | " | Data3(Largest FOV) Fig 2a, Movie 3 | 40x/0.8NA | 40x/0.8NA | NA | 0.33um isotropic | Nematode embryo expressing GCaMP3 | 2.86 volumes per second | 40umx60um | " |
| SCAPE(*11*) | Gaussian | Data1(Highest Spatial Resolution) Sup Fig. 6 | 20x/1.0NA | Same as Excitation Objective | 2.5xumx2umx4um | Lateral: 0.4-2um Axial: 1-3um | 200nm beads | NA | 1000umx550um (y' by z') | Demonstrated usable range of approximately 550 μm. The optimal focal performance occurs 200–400 μm below the sample surface, with diagonal PSF stretching present at the surface and in deeper regions. |
| " | " | Data2(Fastest Temporal Resolution) Sup Fig. 10 | 20x/1.0NA | Same as Excitation Objective | 2.5xumx2umx4um | Lateral: 0.4-2um Axial: 1-3um | First instar myosin heavy-chain *mhc-Gal4,UAS-CD8:GFP* larvae | 20 volumes per second | 1330umx134um (y' by z') | " |
| " | " | Data3(Largest FOV) Fig. 2d | 20x/1.0NA | Same as Excitation Objective | 2.5xumx2umx4um | Lateral: 0.4-2um Axial: 1-3um | Mouse expressing GCaMP5g in layer 5 pyramidal neurons | 10 volumes per second | 650umx134um (y' by z') | " |
| Airy Light Sheet(*12*) | Airy | Data1(Highest Spatial Resolution) Fig. 2,6 | 40x/0.8NA | 40x/0.8NA | Lateral: 1.18um Axial: 1.44um (At surface) | NA | Cleared mouse brain tissue injected with red fluorescent microspheres (diameter 600 nm) | NA | ~332umx332um (Inferred) | ALSM resolves bead size and position at depths of 330 ± 1 μm. |
| " | " | Data2(Fastest Temporal Resolution) Fig. 2,6 | 40x/0.8NA | 40x/0.8NA | Lateral: 1.18um Axial: 1.44um (At surface) | NA | Cleared mouse brain tissue injected with red fluorescent microspheres (diameter 600 nm) | NA | ~332umx332um (Inferred) | " |
| " | " | Data3(Largest FOV) Fig. 2,6 | 40x/0.8NA | 40x/0.8NA | Lateral: 1.18um Axial: 1.44um (At surface) | NA | Cleared mouse brain tissue injected with red fluorescent microspheres (diameter 600 nm) | NA | ~332umx332um (Inferred) | " |
| meSPIM(*13*) | Bessel | Data1(Highest Spatial Resolution) Fig 1K | 40x/0.8NA | 40x/0.8NA | 300-340nm isotropic | 300-340nm isotropic | 100nm fluorescent beads | NA | 163umux163um | Maintains near-isotropic resolution (300–340 nm) and uniform illumination across a 100 μm propagation range. |
| " | " | Data2(Temporal Resolution) Fig 5 | 40x/0.8NA | 40x/0.8NA | 300-340nm isotropic | 300-340nm isotropic | Tracking of membrane blebs on MV3 cells | 1.2 seconds per volume | 100umx100um | " |
| " | " | Data3 (Largest FOV ) Fig 1K | 40x/0.8NA | 40x/0.8NA | 300-340nm isotropic | 300-340nm isotropic | 100nm fluorescent beads | NA | 163umux163um | " |
| SOPi(*14*) | Gaussian | Data1(Highest Spatial Resolution) Fig. 5c | 20x/1.0NA | Same as Excitation Objective | 1P Lateral: 1.3um 2P Lateral: 1.16um | NA | Beads | NA | NA | NA |
| " | " | Data2(Fastest Temporal Resolution) Fig. 7 | 20x/1.0NA | Same as Excitation Objective | 1P Lateral: 1.3um 2P Lateral: 1.16um | NA | GCaMP6s-expressing zebrafish larvae | 10 volumes per second | 850 × 300 × 50 $\mu m^3$ | NA |
| " | " | Data3(Largest FOV) Fig. 6 | 20x/1.0NA | Same as Excitation Objective | 1P Lateral: 1.3um 2P Lateral: 1.16um | NA | Fixed, not optically cleared 1 mm thick section of Thy1-GFP | 30 seconds per volume | 750 × 270 × 500 $\mu m^3$ | NA |

| | | | | | | | | | | |
|---|---|---|---|---|---|---|---|---|---|---|
| | | | | | | | transgenic mouse hippocampus | | | |
| Bessel CBS(*15*) | Bessel | Data1(Highest Spatial Resolution) Fig. 1 | 40x/0.8NA | 40x/0.8NA | Taken but not quantified. | Axial: 1.07um | According to experimental beam profile. | NA | 300 µm × 200 µm × 100 µm | Bessel light-sheets expand FOV and signal but cause axial elongation. Conversely, the CBS method matches Gaussian axial resolution at the center and margins of the FOV while maintaining the uniform illumination characteristic of Bessel beams. |
| " | " | Data2(Fastest Temporal Resolution) Fig. 6 | 40x/0.8NA | 40x/0.8NA | NA | Axial: 1.07um | eGFP-labeled mouse brain section | 40 seconds per volume | 300 µm × 200 µm × 200 µm | " |
| " | " | Data3(Largest FOV) Fig. 6 | 40x/0.8NA | 40x/0.8NA | NA | Axial: 1.07um | eGFP-labeled mouse brain section | 40 seconds per volume | 300 µm × 200 µm × 200 µm | " |
| SCAPE 2.0(*16*) | Gaussian | Data1(Highest Spatial Resolution) Supp Note 1 | 20x / 1.0 NA | 20x / 1.0 NA<br>10x/0.45 (O3) | X:1.47um<br>Y:0.86um<br>Z:1.96um | NA | 200 nm beads | NA | NA | NA |
| " | " | Data2(Fastest Temporal Resolution) Fig 3b | 20x / 1.0 NA | 20x / 1.0 NA<br>20x/ 0.6 (O3) | NA | Cellular resolution | 3 days post-fertilization (d.p.f.) zebrafish embryos expressing GFP and DsRed in its endothelial cells and RBCs | 321 volumes per second | 179umx342umx127um | NA |
| " | " | Data3(Largest FOV) Fig 4e | 20x / 1.0 NA | 20x / 1.0 NA<br>10x/0.45 (O3) | NA | NA | Coronal hemi-section of an mCUBIC-cleared *Thy1*-GFP brain using stage-scanning and stitching | ~4.03 min per volume | ~8.5 × 9.5 × 0.46 $mm^3$ | Used low NA light sheets to span a depth range of up to ~450µm.<br><br>The galvo was swept over a range of 500 µm with a camera frame acquired every 0.5 µm. |
| Mesolens(*17*) | Airy | Data1(Highest Spatial Resolution) Fig. 6 | NA | 4x/0.47NA | NA | Sub-cellular | Cleared coronal section of the mouse forebrain | 8h per volume | 4.4 mm×3.0 mm | The intensity profile along y allowed FWHM measurement of Gaussian light-sheet thickness as 36.8 mm ± 4.1 mm at y = 0 mm position, 30.1 mm ± 5.3 mm at y = 1.5 mm , and 31.3 mm ± 2.1 mm at y = 3 mm.<br><br>Airy light-sheet intensity profile along y was measured by taking the FWHM of the first maximum only. The thickness of the Airy light-sheet was measured as 7.8 mm ± 1.3 mm at y = 0 mm position, 10.1 mm ± 1.5 mm at y = 1.5 mm , and 11.1 mm ± 0.9 mm at y = 3 mm. |
| " | Gaussian | Data2(Fastest Temporal Resolution) Fig. 4 | NA | 4x/0.47NA | NA | NA | Mouse pancreas cleared in a 1:2 mixture of benzyl alcohol:benzyl benzoate (BABB), with nuclei stained with propidium iodide | 4h per volume | 4.4 mm×3.0 mm | " |
| " | Gaussian | Data3(Largest FOV) Fig 4. | NA | 4x/0.47NA | NA | NA | Mouse pancreas cleared in a 1:2 mixture of benzyl alcohol:benzyl benzoate (BABB), with | 4h per volume | 4.4 mm×3.0 mm | " |

| | | | | | | | | | | |
|---|---|---|---|---|---|---|---|---|---|---|
| | | | | | | | nuclei stained with propidium iodide | | | |
| DaXi(*18*) | Gaussian | Data1(Highest Spatial Resolution) Fig. 2 | 20x/1.0NA | Custom AMS-AGY v2.0 (1.0 NA) | Lateral: 0.45um<br>Axial: 2um | NA | 100nm fluorescent beads | NA | 3,000 × 800 × 300 μm | Performance remains uniform across all three channels up to a depth of approximately 300 μm. |
| " | " | Data2(Fastest Temporal Resolution) Extended data fig. 8 | 20x/1.0NA | Custom AMS-AGY v2.0 (1.0 NA) | " | NA | Whole-brain, neuron-level 3D imaging in larval zebrafish | 3.3 volumes per second | 500umx300umx200um | " |
| " | " | Data3(Largest FOV) Fig. 3a | 20x/1.0NA | Custom AMS-AGY v2.0 (1.0 NA) | " | NA | *D. rerio* (zebrafish) larvae at 30 hpf | 52 seconds per volume | 3,000 × 800 × 300 μm | " |
| RUSH3D(*19*) | Scanning Light-field | Data1(Highest Spatial Resolution) Fig. S3A | 2x/0.5 NA | 2x/0.5 NA | Lateral: 2.6um<br>Axial: 6um | Single cell resolution | 500nm beads | NA | NA | RUSH3D maintains an effective depth of field up to 400 μm, defined as the range where lateral and axial resolutions remain within twice their optimal values. |
| " | " | Data2(Fastest Temporal Resolution) Fig. 3 | 2x/0.5 NA | 2x/0.5 NA | Lateral: 2.6um<br>Axial: 6um | Single cell resolution | Head-fixed imaging on layer-2/3-specific transgenic mice with GCaMP6f labeling across 17 cortical regions | 20 volumes per second | 8,000 × 6,000 × 400 $\mu m^3$ | " |
| " | " | Data3(Largest FOV) Fig. 3 | 2x/0.5 NA | 2x/0.5 NA | Lateral: 2.6um<br>Axial: 6um | Single cell resolution | Head-fixed imaging on layer-2/3-specific transgenic mice with GCaMP6f labeling across 17 cortical regions | 20 volumes per second | 8,000 × 6,000 × 400 $\mu m^3$ | " |
| miOPM (20) | Gaussian | Data1(Highest Spatial Resolution) Fig. 5b | 40x/1.33NA | 40x/1.33NA AMS-AGY v2 (O3) | X: 358.1±12.5nm<br>Y: 316.9±11.2nm<br>Z: 753.6±40.9nm | Subcellular | 100 nm nanospheres in Agarose | NA | 56umx100umx110um | NA |
| " | " | Data2(Fastest Temporal Resolution) Fig. 2b | 40x/1.33NA | 40x/1.33NA AMS-AGY v2 (O3) | NA | Subcellular | CD8+ T-cells (membrane labeled with CellMask Orange) | 1s volume rate | 217umx236umx96um | NA |
| " | " | Data3(Largest FOV) Fig. 3 | 40x/1.33NA | 40x/1.33NA AMS-AGY v2 (O3) | NA | Subcellular | 4x expanded liver slice (labelled for chromatin/nucleic acid) | NA | 240umx217umx597um | Vertical height of ~600 μm achieved via vertical tiling of eleven individual volumes. |

### Supplementary table T2

| **Figure/Movie** | **Specimen** | **Fluorophores** | **Airy strength** | **Exposure (Laser)** | **Detection objective** | **Pixel size** | **Step size** | **Image stack size (XYZ)** | **Volume of imaging (XYZ)** | **Time per volume** |
|---|---|---|---|---|---|---|---|---|---|---|
| 1f–n; 2b,c | zebrafish, wt | phalloidin-Alexa Fluor 647 | 5, 30, 90 | 50 ms (647) | 25× 1.1 NA | 130 nm | 200 nm | 2048 × 2048 × 1600 $px^3$ | 266 × 266 × 320 $\mu m^3$ | 93.1 s |
| 2a | RPE1, ER-StayGold Mito-HaLo | JFX650 | 5, 30, 90 | 10 ms (647) | 20× 1.0 NA | 162 nm | 200 nm | 2048 × 2048 × 1001 $px^3$ | 332 × 332 × 200 $\mu m^3$ | 18.2 s |
| 3a–e, S4a,b; SM2; SM3; SM4 | zebrafish tail 3 dpf, cross of *Tg(actc1b:mito-GFP)*$^{uom407Tg}$ and *Tg(actc1b:ER-mCherry)*$^{uom408Tg}$ | mito-GFP, ER-mCherry | 70 | 30 ms (488), 30 ms (561) | 25× 1.1 NA | 130 nm | 200 nm | 2048 × 2048 × 200 $px^3$ | 266 × 266 × 40 $\mu m^3$ | 19.0 s |
| 4a–e; SM5; SM6 | zebrafish tailbud (14 somite stage), *Tg(fgf8a:fgf8a-EGFP)* injected with mRNA for mKate2-rab5ab | mKate2-rab5ab | 70 | 20 ms (561) | 25× 1.1 NA | 130 nm | 400 nm | 2048 × 2048 × 151 $px^3$ | 266 × 266 × 60 $\mu m^3$ | 5.9 s |
| 4f–h | zebrafish tail 1 dpf, wt injected with mRNA for mKate2-rab5ab and mem-mNeonGreen | membrane-mNeonGreen, mKate2-rab5ab | 70 | 50 ms (488), 50 ms (561) | 25× 1.1 NA | 130 nm | 200 nm | 2048 × 2048 × 90 $px^3$ | 266 × 266 × 18 $\mu m^3$ | 15.3 s |
| 5a–g; S6a,b; SM7; SM8; SM9; SM10 | *Drosophila* embryo, sqh-3x-GFP | sqh-3x-GFP | 50 | 20 ms (488) | 25× 1.1 NA | 130 nm | 200 nm | 2048 × 2048 × 926 $px^3$ | 266 × 266 × 185 $\mu m^3$ | 24.5 s |
| 6a,b; S8a–d; SM13 | meiosis II oocytes from 7-week-old PhAM mice | mito-Dendra2 | 30 | 20 ms (488) | 25× 1.1 NA | 130 nm | 200 nm | 756 × 728 × 501 $px^3$ | 98 × 94 × 100 $\mu m^3$ | 41.4 s |
| 6c; S9; SM14 | *Amoeba proteus* | DiI | 50 | 10 ms (561) | 20× 1.0 NA | 162 nm | 1000 nm | 1320 × 1644 × 201 $px^3$ | 214 × 267 × 201 $\mu m^3$ | 6.2 s |
| 6d | *Amoeba proteus* | DiI | 70 | 30 ms (561) | 20× 1.0 NA | 162 nm | 1000 nm | 2048 × 2048 × 101 $px^3$ | 332 × 332 × 101 $\mu m^3$ | 5.8 s |
| 7a–c | patient-derived colorectal cancer organoid, TUBB::TagGFP2 | PKmitoDeepRed | 30 | 100 ms (647) | 20× 1.0 NA | 162 nm | 200 nm | 2048 × 2048 × 601 $px^3$ | 332 × 332 × 120 $\mu m^3$ | 64.6 s |
| 7d–f; SM15 | iBlastoid | DAPI, α-NANOG in 488, α-GATA3 in 555, α-GATA6 in 647, phalloidin-Alexa Fluor 680 | 70 | 100 ms (405, 488, 561, 647), 200 ms (685) | 20× 1.0 NA | 162 nm | 200 nm | 2048 × 2048 × 900 $px^3$ | 332 × 332 × 180 $\mu m^3$ | 576.9 s |
| S3b,c; SM1 | HeLa, Rab5-GFP Rab7-mCherry | Rab5-GFP, Rab7-mCherry | 5 | 20 ms (488+561) | 20× 1.0 NA | 162 nm | 200 nm | 636 × 1864 × 126 $px^3$ | 103 × 302 × 25 $\mu m^3$ | 5.7 s |
| S5b,cA | zebrafish tail 3 dpf, *Tg(acta1:mCherryCAAX)*$^{pc22Tg}$ | CaaX-mCherry | 70 | 30 ms (561) | 25× 1.1 NA | 130 nm | 200 nm | 2048 × 2048 × 763 $px^3$ | 266 × 266 × 153 $\mu m^3$ | 150.0 s |
| S7a–c; SM11 | *Drosophila* embryo, sqh-3x-GFP | sqh-3x-GFP | 70 | 30 ms (488) | 25× 1.1 NA | 130 nm | 200 nm | 2048 × 3864 × 400 $px^3$ | 266 × 501 × 80 $\mu m^3$ | 66.0 s |
| SM12 | *Drosophila* embryo, sqh-3x-GFP | sqh-3x-GFP | 70 | 50 ms (488) | 25× 1.1 NA | 130 nm | 200 nm | 2048 × 2048 × 688 $px^3$ | 266 × 266 × 138 $\mu m^3$ | 120.0 s |

#### References


1. J. Huisken, J. Swoger, F. Del Bene, J. Wittbrodt, E. H. Stelzer, Optical sectioning deep inside live embryos by selective plane illumination microscopy. *Science* **305**, 1007-1009 (2004).
2. P. J. Keller *et al.*, Fast, high-contrast imaging of animal development with scanned light sheet-based structured-illumination microscopy. *Nat Methods* **7**, 637-642 (2010).
3. T. A. Planchon *et al.*, Rapid three-dimensional isotropic imaging of living cells using Bessel beam plane illumination. *Nature Methods* **8**, 417-423 (2011).
4. U. Krzic, S. Gunther, T. E. Saunders, S. J. Streichan, L. Hufnagel, Multiview light-sheet microscope for rapid in toto imaging. *Nat Methods* **9**, 730-733 (2012).
5. R. Tomer, K. Khairy, F. Amat, P. J. Keller, Quantitative high-speed imaging of entire developing embryos with simultaneous multiview light-sheet microscopy. *Nat Methods* **9**, 755-763 (2012).
6. B. C. Chen *et al.*, Lattice light-sheet microscopy: imaging molecules to embryos at high spatiotemporal resolution. *Science* **346**, 1257998 (2014).
7. T. Vettenburg *et al.*, Light-sheet microscopy using an Airy beam. *Nature Methods* **11**, 541-544 (2014).
8. R. K. Chhetri *et al.*, Whole-animal functional and developmental imaging with isotropic spatial resolution. *Nat Methods* **12**, 1171-1178 (2015).
9. T. L. Liu *et al.*, Observing the cell in its native state: Imaging subcellular dynamics in multicellular organisms. *Science* **360**, (2018).
10. Y. Wu, *et al.*, Reflective imaging improves spatiotemporal resolution and collection efficiency in light sheet microscopy. *Nature Communications* **8**, 1452 (2017).
11. M. B. Bouchard *et al.*, Swept confocally-aligned planar excitation (SCAPE) microscopy for high-speed volumetric imaging of behaving organisms. *Nature Photonics* **9**, 113-119 (2015).
12. J. Nylk, K. McCluskey, S. Aggarwal, J. A. Tello, K. Dholakia, Enhancement of image quality and imaging depth with Airy light-sheet microscopy in cleared and non-cleared neural tissue. *Biomed Opt Express* **7**, 4021-4033 (2016).
13. E. S. Welf *et al.*, Quantitative Multiscale Cell Imaging in Controlled 3D Microenvironments. *Dev Cell* **36**, 462-475 (2016).
14. M. Kumar, S. Kishore, J. Nasenbeny, D. L. McLean, Y. Kozorovitskiy, Integrated one- and two-photon scanned oblique plane illumination (SOPi) microscopy for rapid volumetric imaging. *Opt. Express* **26**, 13027-13041 (2018).
15. H. Jia *et al.*, Axial resolution enhancement of light-sheet microscopy by double scanning of Bessel beam and its complementary beam. *J Biophotonics* **12**, e201800094 (2019).
16. V. Voleti *et al.*, Real-time volumetric microscopy of in vivo dynamics and large-scale samples with SCAPE 2.0. *Nature Methods* **16**, 1054-1062 (2019).
17. E. Battistella, J. Schniete, K. Wesencraft, J. F. Quintana, G. McConnell, Light-sheet mesoscopy with the Mesolens provides fast sub-cellular resolution imaging throughout large tissue volumes. *iScience* **25**, 104797 (2022).
18. B. Yang *et al.*, DaXi—high-resolution, large imaging volume and multi-view single-objective light-sheet microscopy. *Nature Methods* **19**, 461-469 (2022).
19. Y. Zhang *et al.*, Long-term mesoscale imaging of 3D intercellular dynamics across a mammalian organ. *Cell* **187**, 6104-6122.e6125 (2024).
20. B. Chen *et al.*, Multi-immersion Oblique Plane Microscope (miOPM): A reconfigurable platform for high-resolution Light-Sheet Fluorescence Microscopy. *bioRxiv*, (2025).
